# Myogenic and neural transcriptional circuitry determines PAX3::FOXO1-driven initiation of rhabdomyosarcoma

**DOI:** 10.64898/2026.10.27.761987

**Authors:** Bradley T. Stevens, Yang Zhang, Matthew R. Garcia, Jack D. Hopkins, Frank S. Wilbanks, Muna N. Ogwo, Shaina N. Porter, Brooke B. Carter, Edwin D. Mathew, Kyna Vuong, Hanna Gebremichael, Uma Neelakantan, Sashi Mandava, Randolph K. Larsen, Francesca Ferrara, Robert E. Throm, Anoop M. Kavirayani, Shondra M. Pruett-Miller, Brian J. Abraham, Mark E. Hatley

**Affiliations:** Department of Oncology, St. Jude Children’s Research Hospital, Memphis, TN, 38105, USA; St. Jude Graduate School of Biomedical Sciences, Memphis, TN, 38105, USA; Department of Computational Biology, St. Jude Children’s Research Hospital, Memphis, TN, 38105, USA; Department of Cell and Molecular Biology and Center for Advanced Genome Engineering, St. Jude Children’s Research Hospital, Memphis, TN, 38105, USA; St. Jude Vector Laboratory, St. Jude Children’s Research Hospital, Memphis, TN, 38105, USA; Department of Pathology, St. Jude Children’s Research Hospital, Memphis, TN, 38105, USA

**Keywords:** rhabdomyosarcoma, PAX3::FOXO1, sarcoma, fusion oncoprotein, oncogenic transcription factor

## Abstract

Alveolar rhabdomyosarcoma (ARMS) is largely characterized by the PAX3::FOXO1 (P3F) fusion oncoprotein, yet the mechanisms by which P3F initiates tumorigenesis and cell identity remain incompletely understood. Here, we describe a doxycycline-inducible, degradable PAX3::FOXO1 iPSC-derived ARMS model (ddP3F) to interrogate the temporal requirements of P3F for ARMS cell identity and proliferation. We show that, while P3F is essential for initiating ARMS transformation from precursor cells, continued expression of P3F is not required for maintaining proliferation, viability, or RMS-associated transcriptional programs. Integration of CUT&RUN, bulk RNA-seq, and 10X multiome profiling during endothelial-directed differentiation revealed that P3F rapidly pioneers a novel enhancer landscape within two days of expression, activating transcriptional circuits anchored by both specific myogenic and neural transcription factors (TFs), including MYOD1, PRDM12, and POU4F1 (BRN3A). Our single-cell analysis shows that these TFs are specifically co-expressed with P3F from the earliest stages of transformation. Knockout of MYOD1 or POU4F1 impairs ARMS identity acquisition without preventing transformation, whereas PRDM12 loss enhances P3F expression programs and myogenic differentiation. Together, these findings underscore a previously unappreciated interplay between myogenic and neural transcriptional programs during ARMS initiation. Furthermore, these data demonstrate that P3F establishes a stable oncogenic state that persists even after removal, thus providing a framework for evaluating P3F-dependent and P3F-independent therapeutic vulnerabilities.

**STATEMENT OF SIGNIFICANCE:** This study reveals that PAX3::FOXO1 initiates rhabdomyosarcoma by establishing myogenic and neural transcriptional programs, yet becomes dispensable for tumor maintenance, uncovering stable oncogenic states and highlighting opportunities for targeting early transforming events.

## INTRODUCTION

Rhabdomyosarcoma (RMS) is the most common pediatric soft tissue sarcoma, and despite numerous efforts, the survival rate for patients with high-risk RMS has remained at 20% for decades^1–3^. The two major subtypes of RMS are alveolar RMS (ARMS) and embryonal RMS (ERMS), defined by histological features and genetic drivers^4, 5^. The majority of ARMS tumors harbor either t(2;13)(q35;q14) or t(1;13)(q36;q14) chromosomal translocations that result in PAX3/7::FOXO1 (P3F and P7F respectively) fusion oncoproteins. Children with ARMS that lack these translocations have tumors with gene expression profiles and survival indistinguishable from ERMS. Therefore, RMS is more accurately molecularly classified as fusion-positive RMS (FP-RMS) for those harboring the PAX3/7::FOXO1 oncofusion proteins, and fusion-negative RMS for those that do not^6–9^. FP-RMS tumors have low mutational burden, averaging only 6.4 mutations per genome, suggesting that fusion oncoproteins drive the disease^10–12^.

P3F is a chimeric transcription factor (TF) that results from fusing the DNA-binding domains of PAX3 and the transactivation domain of FOXO1. P3F acts as a strong activating TF that has been shown to bind heterochromatin and induce enhancers through direct interactions with the histone acetyltransferase EP300 and members of the SWI/SNF chromatin remodeling complex^13–16^. These activities are believed to initiate RMS cell identity and transformation; however, the specific targets and processes governing ARMS tumor initiation remain unknown. RMS tumor cells resemble skeletal muscle progenitor cells that have failed to terminally differentiate leading to the theory that skeletal muscle progenitors are the cell-of-origin for RMS tumors. However, other cells-of-origin have been identified previously for ARMS, including endothelial cells, suggesting a unified mechanism of action for the fusion oncoprotein in transformation and identity acquisition from multiple putative origin points^17, 18^.

Previously, the Hatley lab generated an in vitro model system whereby human induced pluripotent stem cell (iPSC)-derived ARMS (iARMS) cells are generated through forced expression of P3F during endothelial cell directed differentiation^17^. The iARMS system provides multiple opportunities for understanding the mechanism of action of P3F in ARMS initiation in human cells by (i) allowing access to key timepoints during differentiation and transformation, (ii) from a non-myogenic cell of origin, (iii) with no cooperating transforming mutations required. Previous attempts to generate ARMS cells from stem cells either required multiple genetic manipulations prior to P3F expression or forced expression of muscle specification transcription factors prior to P3F addition^19, 20^. Given that the iARMS system originates from human progenitor cells not destined to become muscle cells, probing timepoints throughout iARMS differentiation allows for understanding of the sequence of changes orchestrated by P3F to define the key determinants of iARMS cell identity in a human model system.

Here, we adapted our iARMS system through transduction of pLenti-TRE3G-P3F-FKBP12^F36V^-HA, encoding a degron fusion, into BJFF.6 iPSCs to directly test and probe maintenance of iARMS cell proliferation and identity upon P3F degradation. We show that, upon degradation of P3F, iARMS cells remain viable, continue to proliferate, do not significantly differentiate, and maintain RMS cell identity. We further show the order of operations of P3F-mediated transformation and establishment of RMS cell identity through epigenetic re-wiring of endothelial progenitor cells concomitant with expression of myogenic and neural TFs. We identify a set of early-response TFs upregulated during transformation, and find that deletion of these key TFs alters acquisition of RMS identity and transcriptional circuitry. Taken together, our data reveal that P3F quickly initiates a myogenic transcriptional program in collaboration with neural TFs to establish RMS identity.

## RESULTS

### Generation of doxycycline-inducible, degradable P3F human iPSC model

Analysis of the Cancer Dependency Map indicates that both *PAX3* and *FOXO1* are genetic dependencies in ARMS cells (Figure 1a-c)^21^. This observation suggests that targeting P3F could have therapeutic efficacy; however, interfering with P3F remains to be rigorously tested using in vivo models. Furthermore, genetic deletion is a non-physiologic representation of depleting a gene of interest. Analysis of the DEMETER shRNA screening data revealed no clear dependency with knockdown of either *PAX3* or *FOXO1* (Figure 1d-f)^21^. Additionally, P3F knockdown with siRNA/shRNA has shown variable effects on focus-formation, motility, and cell viability in a variety of ARMS cell lines^22–26^. Furthermore, targeted degradation of P3F in P3F-FKBP12^F36V^ engineered Rh4 clones showed more consistent decreases in proliferation and viability with increased differentiation, while P3F-FKBP12^F36V^ engineered Rh30 clones remained heterogeneous^14^. Moreover, directly inhibiting P3F activity with piperacetazine decreased anchorage-independent growth of multiple ARMS cell lines but did not alter tumor growth in xenografted cells^27^. The most perplexing observation was that P3F-transformed human myoblasts formed ARMS tumors that regressed upon P3F removal but subsequently recurred without P3F fusion protein expression^28^. These recurring-tumors maintained ARMS-like histomorphology and expression of RMS markers; however, it remains unclear how the cells escaped P3F deletion^28^. Taken together, these data suggest that P3F is important for tumor initiation but may be dispensable in tumor maintenance. Since current standard-of-care therapies do not provide a selective pressure for ARMS cells to lose P3F expression, a better understanding of the initiation mechanisms and maintenance dependence are needed prior to advancing P3F-directed therapy.

**Figure 1.**
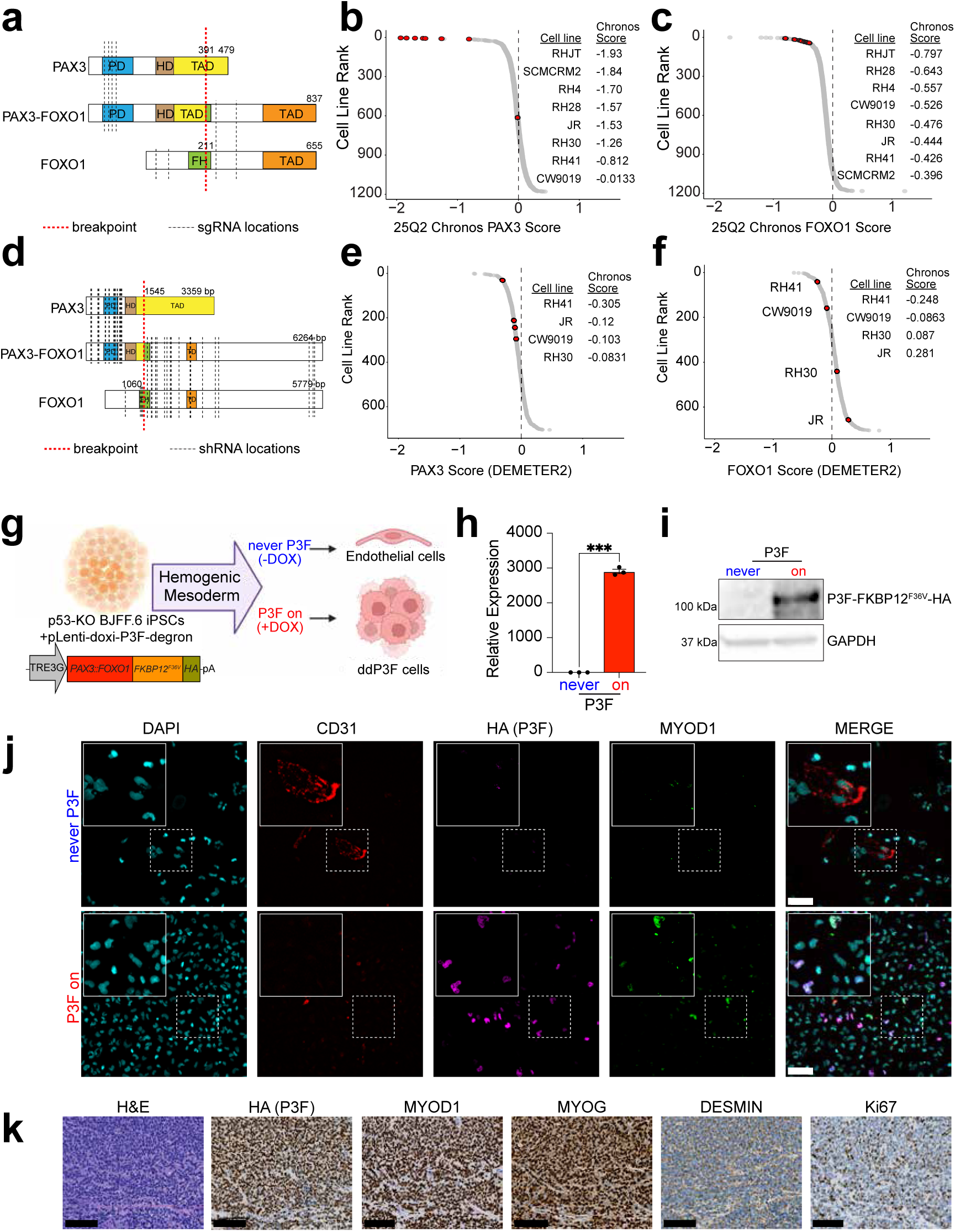
Generation of doxycycline-inducible, degradable PAX3::FOXO1 (ddP3F) human iPSC model. **a)** Schematic of PAX3::FOXO1 oncofusion protein with breakpoint (red dashed line) and sgRNA locations (black dash lines) used in DepMap to target *PAX3*, *FOXO1*, and subsequently *PAX3::FOXO1*. **b)** 25Q2 CHRONOS dependency scores for *PAX3* in all cell lines in DepMap. Highlighted points (red) represent ARMS cell lines, with the individual scores listed. Chronos scores < -0.5 represent candidate as a dependency. **c)** 25Q2 CHRONOS dependency scores for *FOXO1* in all cell lines in DepMap. Highlighted points (red) represent ARMS cell lines, with the individual scores listed. Chronos scores < -0.5 represent candidate as a dependency. **d)** Schematic of *PAX3::FOXO1* oncofusion protein with breakpoint (red dashed line) and shRNA locations (black dash lines) used in shRNA screen from Project Achilles to target *PAX3*, *FOXO1*, and subsequently *PAX3::FOXO1*. **e)** DEMETER2 dependency scores for *PAX3* in all cell lines screened in DepMap. Highlighted points (red) represent ARMS cell lines, with the individual scores listed. **f)** DEMETER2 dependency scores for *FOXO1* in all cell lines in DepMap. Highlighted points (red) represent ARMS cell lines, with the individual scores listed. **g)** Schematic of iPSC directed differentiation generating ddP3F cells. **h)** *PAX3::FOXO1* expression by real-time PCR (n = 3, p < 0.001) from “never P3F” and “P3F on” iPSCs. **i)** Immunoblots of PAX3::FOXO1 and GAPDH on “never P3F” and “P3F on” iPSCs. **j)** Representative immunofluorescence of DAPI (cyan), CD31 (red), HA (magenta), and MYOD1 (green) on “never P3F” and “P3F on” ddP3F cells at Day 15 of endothelial-directed differentiation and ddP3F cell generation. Scale bar = 100 µm. **k)** Representative histology (H&E) and immunohistochemistry (HA (P3F), MYOD1, MYOG, DESMIN, and Ki67) of “P3F on” ddP3F tumors. Scale bar = 100 µm. All p-values in pairwise comparisons were determined by Student’s t-test (unpaired, two-tailed); ***p < 0.001. Data represented as mean +/- SEM.

To further refine our understanding of P3F’s roles in RMS tumor initiation and maintenance, we modified our iARMS system to include a doxycycline-inducible and degradable form of P3F (ddP3F cells). We transduced *TP53* KO BJFF.6 iPSCs with a doxycycline-inducible P3F fused to an FKBP12^F36V^ degron tag and an HA epitope (Figure 1g). This construct allowed for doxycycline-inducible *P3F* expression as well as cessation of *P3F* expression with doxycycline withdrawal. The addition of the FKBP12^F36V^ degron tag allows for P3F degradation with dTAG^V^-1 administration^29, 30^. Culturing cells in media containing 10 µg/mL doxycycline resulted in robust P3F mRNA and protein expression within 24-48 hours (“P3F on”) (Figure 1h-i). Following doxycycline withdrawal, P3F expression is not detected by immunoblot after 4 days (Supplemental Figure 1a). Furthermore, cells treated with 100 nM dTAG^V^-1 degrade P3F nearly completely within 4 hours and do not re-express P3F after doxycycline and dTAG^V^-1 washout (Supplemental Figure 1b-c). This ddP3F system allows for acute control over P3F expression to allow for rigorous testing of P3F targeting.

To build on our previous method for generating iARMS, we induced P3F expression with doxycycline beginning at day 5 of endothelial-directed differentiation^17, 31^. Induced P3F expression blocked endothelial differentiation, shown by decreased CD31 staining, and reprogramming to muscle identity evident by increased MYOD1 staining (Figure 1j and Supplemental Figure 1d). Cells not treated with doxycycline did not express P3F (“never P3F”), executed endothelial differentiation, and expressed CD31 (Figure 1j and Supplemental Figure 1d). When xenografted into immunocompromised mice maintained on doxycycline-containing water, “P3F on” ddP3F cells formed tumors resembling solid-variant ARMS (Figure 1k). Taken together, these data show the generation of a “togglable” system by which ARMS can be generated with doxycycline control over the P3F oncogenic fusion protein expression and P3F protein removal controlled by doxycycline withdrawal and dTAG^V^-1-induced degradation.

### RMS identity is retained following loss of P3F

To mimic the effect of maximally targeting P3F, we removed P3F from ddP3F cells that had completed endothelial directed differentiation by simultaneously withdrawing doxycycline and adding 100 nM dTAG^V^-1 (“P3F off”). We then assessed cell proliferation, cell viability, myogenic differentiation, and endothelial tube formation comparing to ddP3F cells maintained in doxycycline and treated with DMSO vehicle (“P3F on”) (Figure 2a). Interestingly, while degradation of P3F affected cell proliferation, viability remained the same compared to “P3F on” ddP3F cells, indicating overall growth was only partially suppressed (Figure 2b-c). Analysis of Incucyte images acquired at day 7 of the cellular proliferation assay indicated that “P3F on” ddP3F cells were more confluent with larger cells and less intercellular space while the “P3F off” ddP3F cells had subtly more intervening space between cells (Supplemental Figure 1e).

**Figure 2.**
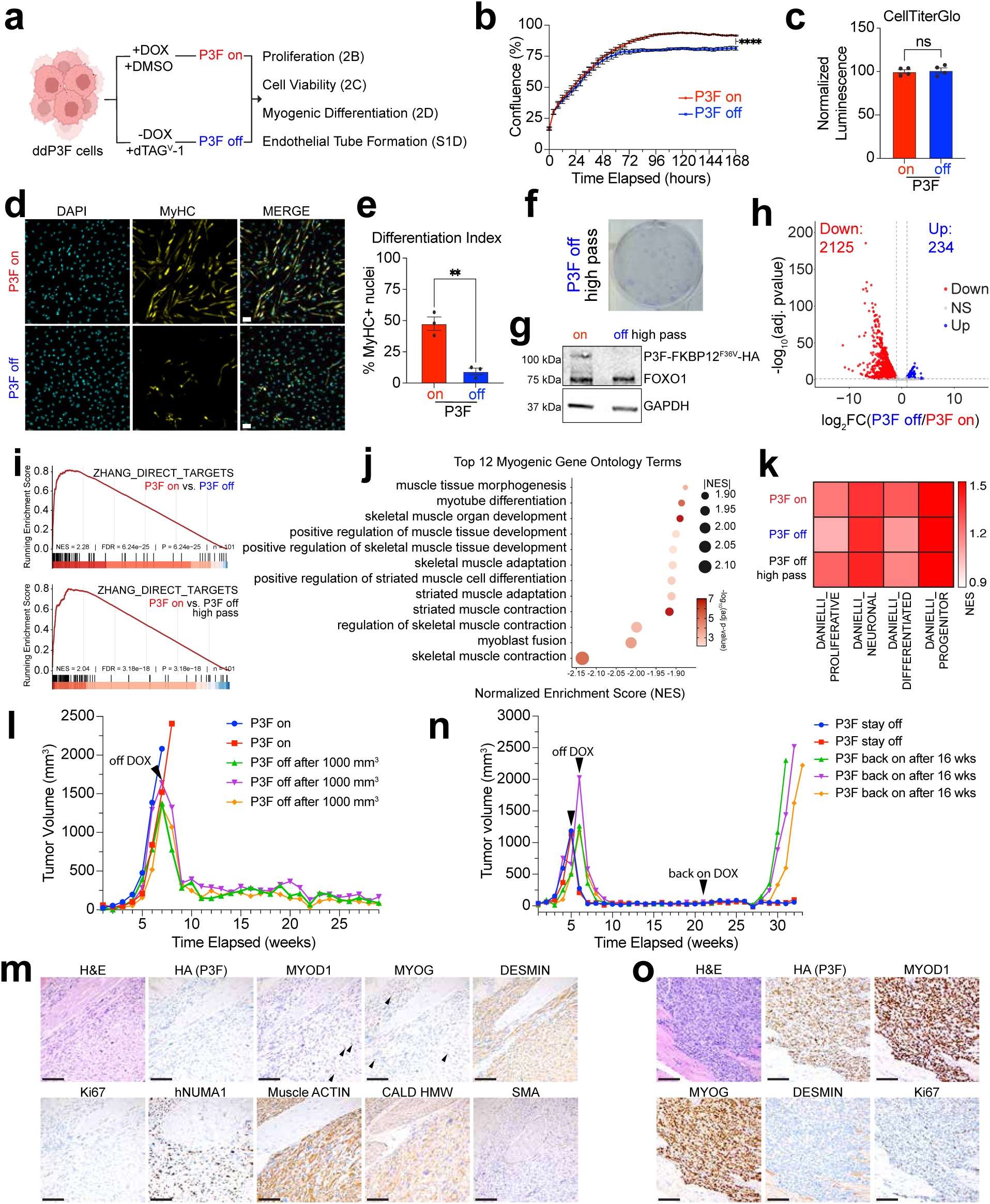
RMS identity is retained following loss of P3F. **a)** Schematic of experiment to generate “P3F off” ddP3F cells through doxycycline removal and dTAG^V^-1 mediated P3F degradation. **b)** Representative cell proliferation data showing confluence over time for both “P3F on” and “P3F off” ddP3F cells. **c)** Representative cell viability data measured via CellTiterGlo assay for both “P3F on” and “P3F off” ddP3F cells. **d)** Representative immunofluorescence of “P3F on” and “P3F off” ddP3F cells for DAPI (cyan) and MyHC (yellow) after culturing for 7 days in myogenic differentiation media. **e)** Differentiation index calculation of “P3F on” and “P3F off” ddP3F cells cultured for 7 days in myogenic differentiation media. n = 3 biological replicates. **f)** Representative well of focus-formation assay on high-passage “P3F off” ddP3F cells. n = 3 wells. **g)** Immunoblot for PAX3::FOXO1 and GAPDH in “P3F on” and “P3F off high pass” ddP3F cells. **h)** Volcano plot showing differentially expressed genes with absolute log_2_FC greater than 1 and adjusted p-value less than 5% in “P3F on” and “P3F off” ddP3F cells. **i)** Enrichment plot of P3F-direct target gene set (ZHANG_DIRECT_P3F_TARGETS identified by Zhang et al.^14^) for “P3F off” and “P3F off high pass” ddP3F cells compared to “P3F on” ddP3F cells. **j)** Dot plot of the top 12 myogenic gene ontology terms enriched in “P3F on” ddP3F cells compared to “P3F off” ddP3F cells. **k)** Heatmap of the gene expression of “P3F on,” “P3F off,” and “P3F off high pass” ddP3F cells. We performed gene set enrichment analysis (GSEA) using a preranked approach for analyses not based on direct condition-to-condition comparisons to score proliferative (DANIELLI_PROLIFERATIVE), neuronal (DANIELLI_NEURONAL), differentiated (DANIELLI_DIFFERENTIATED), and progenitor (DANIELLI_PROGENITOR) cell states identified by Danielli et al.^35^. Genes were ranked by expression (TPM) rather than differential expression. **l)** Tumor volume measurements for NSG mice injected with “P3F on” ddP3F cells and kept on doxycycline containing water. “P3F off after 1000 mm^3^” tumors were generated by replacing mice into cages with non-doxycycline containing water. Arrowhead indicates when mice were removed from doxycycline treatment. **m)** Representative histology (H&E) and immunochemistry (HA (P3F), MYOD1, MYOG, DESMIN, Ki67, hNUMA1, muscle ACTIN, Caldesmon HMW, and SMA) of “P3F off after 1000 mm^3^” tumor displaying characteristic features of pleiomorphic leiomyosarcoma. **n)** Tumor volume measurements for NSG mice injected with “P3F on” ddP3F cells, kept on doxycycline containing water, and then removed once tumors reached 1000 mm^3^. “P3F back on after 16 wks” tumors were generated by returning mice to treatment with doxycycline containing water 16 weeks after being removed from treatment with doxycycline containing water. “P3F stay off” mice were generated by maintaining the mice on non-doxycycline containing water. **o)** Representative histology (H&E) and immunohistochemistry (HA(P3F), MYOD1, MYOG, DESMIN, and Ki67) of “P3F back on after 16 wks” tumors displaying characteristic features of solid-type ARMS.

Degradation of P3F decreased myogenic differentiation, illustrated by a reduction in the number of cells expressing myosin heavy chain (Figure 2d-e). Remarkably, ddP3F cells continued to proliferate without P3F for greater than 20 passages (“P3F off high pass”), approximately 5 months, and formed colonies in a focus-formation assay while maintaining no expression of P3F (Figure 2f-g). Both “P3F on” and “P3F off” ddP3F cells formed endothelial-like tube structures similar to human umbilical vein endothelial cells (HUVEC) and in contrast to immortalized human myoblasts that failed to form tubes (Supplemental Figure 1f). This finding that ddP3F cells retain the capacity to form endothelial-like tube structures suggests the retention of features of the cell of origin. To test whether this phenomenon was observed in human-derived ARMS cell lines, we performed endothelial tube forming assays with human ARMS cell lines. Rh30 and JR cells form endothelial-like tube structures, while Rh3, Rh4, and Rh41 cells rarely formed tube-like structures (Supplemental Figure 1f-g). Among the ARMS cell lines that retained the ability to form endothelial-like tube structures, similar numbers and sizes of tube structures were observed (Supplemental Figure 1g-h). Thus, the ability of RMS-like cells to form tube structures is a common but not universal feature that may relate to cells of origin.

Initially, we hypothesized that iARMS cells required P3F expression to maintain ARMS cell identity and survival. However, the observation that ddP3F cells lacking P3F continued to proliferate and formed colonies in a focus-formation assay suggested otherwise. To gain insight into cell identity changes after P3F removal, we performed RNA-seq in established ddP3F cells following 1 week without P3F (“P3F off”). “P3F off” ddP3F cells showed selective changes in gene expression compared to “P3F on” ddP3F cells, predominately downregulation (Figure 2h). Indeed, when compared to “P3F on” ddP3F cells, both “P3F off” and “P3F off high pass” ddP3F cells showed downregulation of direct P3F target genes characterized in the literature (ZHANG_DIRECT_TARGETS, Figure 2i)^14^. Additionally, consistent with the reduction in myogenic differentiation, we observed decreased enrichment for multiple gene ontology terms related to myogenic differentiation in “P3F off” ddP3F cells compared to “P3F on” ddP3F cells (Figure 2j). While several hundred genes were significantly downregulated, overall expression of RMS cell identity signatures that were curated through multiple single-cell sequencing efforts of patient tumors and cell lines was collectively high in all settings^32–35^. We observed enrichment of each cell identity gene set (DANIELLI_PROLIFERATIVE, DANIELLI_NEURONAL, DANIELLI_DIFFERENTIATED, and DANIELLI_PROGENITOR) in each of “P3F-on”, “P3F off”, and “P3F off high pass” ddP3F cells (Figure 2k). Importantly, we saw elevated expression of these signature genes without re-expression of P3F (Supplemental Figure 1i). Together, these data suggest that changes in cell identity-related gene expression upon P3F loss might be subtle, and that P3F may not be required for maintaining their expression.

Since our ddP3F model allows for both rapid and slow removal of P3F, we were curious if the rate of P3F loss resulted in similar transcriptional changes. To test this, we cultured ddP3F cells in doxycycline-free media without dTAG^V^-1 for 1 week to gradually turn of P3F expression (“slow off”) and performed RNA-seq. Similar to “P3F off” ddP3F cells, “slow off” ddP3F cells showed changes in transcription mostly associated with downregulation (Supplemental Figure 1j). Directly comparing “P3F off” and “slow off” ddP3F cells revealed virtually no significant differences in transcriptional programs (Supplemental Figure 1k). Furthermore, “slow off” ddP3F cells showed similar changes in gene expression for direct P3F targets (Supplemental Figure 1l). Finally, we observed that “slow off” ddP3F cells still showed enrichment for RMS identity gene sets without re-expression of P3F (Supplemental Figure 1i and 1m). Interestingly, the ARMS cell identity gene sets also showed enrichment in cells from Day 0 and Day 5 of our differentiation protocol (timepoints prior to P3F expression), indicating these gene sets may represent a background signature of stemness and/or mesodermal progenitor that could exist in ARMS cells (Supplemental Figure 1m). This indicates that the rate at which ddP3F cells lose P3F does not alter the resulting transcriptional changes.

With the observation that ARMS cell identities are present in the absence of P3F, we sought to understand the impact of P3F removal on enhancer landscape. To this end, we performed Cleavage Under Target & Release Using Nuclease (CUT&RUN) for H3K27ac in “P3F on,” “P3F off,” and “slow off” ddP3F cells. Consistent with gene expression, we observed changes in “P3F off” ddP3F cell enhancers primarily associated with decreased signal (Supplemental Figure 1n). Additionally, “P3F off” and “slow off” ddP3F enhancer landscapes were highly consistent with each other, further supporting our insight that the rate at which ddP3F cells lose P3F does not alter the resulting changes (Supplemental Figure 1o). Finally, we observed both decreased H3K27ac signal and retention at known P3F target genes MYOD1 and ASS1 respectively (Supplemental Figure 1p-q)^14, 36^. Our data indicate that P3F removal results in epigenetic changes at select, but not all P3F target gene associated enhancers.

One important utility of our ddP3F system is the ability to rigorously test the effect of P3F removal in vivo through doxycycline withdrawal. To determine the impact of maximally targeting P3F in vivo we began by passaging ddP3F cells in NSG mice and subsequently maintained them on doxycycline-containing water. We followed two mice to tumor burden, after 7 weeks (P3F on, Figure 2l). Three mice were taken off doxycycline-containing water after tumor volume reached 1000 mm^3^ (P3F off after 1000 mm^3^, Figure 2l). All three tumors regressed, but tumor volume did not reach 0 mm^3^. Instead, the regressed tumors remained consistently around 100mm^3^-200mm^3^. Two of these mice succumbed to illness unrelated to tumor burden and their tumors were unrecoverable. The third mouse was sacrificed and its tumor-bearing leg investigated. The residual tumor was indeed negative for P3F and was sparsely populated with MYOD1 and MYOG positive cells, some of which were associated with reactive myofiber repair (Figure 2m). Tumor cells featured strong DESMIN positive reactivity, but low Ki67 positive reactivity (Figure 2m). Further immunophenotyping showed expression of muscle-specific ACTIN and higher molecular weight CALDESMON with little expression of smooth muscle ACTIN (SMA). Tumor cells also showed positive reactivity to the human cell identity marker hNUMA1 (Figure 2m). Based on the histopathologic features and immunophenotype, a pathology diagnosis of pleiomorphic leiomyosarcoma was made. Following this observation, we repeated the experiment with five more mice (Figure 2n). All mice in the repeat experiment were taken off doxycycline-containing water after tumors reached 1000 mm^3^ (Figure 2n). To mimic the impact of cessation of P3F targeted therapy, we returned three mice to doxycycline-containing water 16 weeks after doxycycline withdrawal. All three mice succumbed to tumors which were histopathologically and immunophenotypically characterized as solid type alveolar rhabdomyosarcoma (P3F back on after 16 wks, Figure 2n-o). The remaining two mice not returned to doxycycline-containing water were euthanized at a similar time point and their tumor-bearing legs were histologically analyzed (P3F stay off, Figure 2n and Supplemental Figure 1r). Microscopic evaluation of these tissues revealed no obvious residual tumor, but remnant xenograft cells (with hNUMA1 positive reactivity) interspersed with normal mouse myocytes and fibroadipose tissue (Supplemental Figure 1r). These data indicate that P3F removal in vivo could result in alternative sarcomatous malignancies and that P3F-targeting therapies could require lifelong treatment due to the persistence of tumor cells even in the absence of a notable mass. Furthermore, the presence of MYOD1-, MYOG-, and DESMIN-positive cells supports our notion that ARMS cell identities can be maintained in the absence of P3F. All told, these bulk expression analyses, enhancer analyses, and in vivo P3F removal studies in our iARMS model show that P3F is important for tumor initiation and establishment of, but not necessarily identity maintenance, of ARMS identity. Thus, we hypothesized that unraveling the P3F-mediated initiation events would provide mechanistic insight into P3F biology and ARMS identity establishment.

### P3F initiates a tumorigenic transcriptional network

Our published iARMS model and current ddP3F model were generated from male-derived *TP53-*null BJFF.6 iPSCs. However, most ARMS tumors in children are wild-type (WT) for *TP53,* and the disease affects both males and females^37^. To better represent etiology, we first tested if *TP53* deletion was necessary and whether there were sex-based differences in human iPSC sources during P3F-mediated transformation. We repeated the iARMS differentiation protocol in both WT *TP53* male-derived BJFF.6 iPSCs and WT *TP53* female-derived AN1.1 iPSCs. P3F expression in endothelial-directed differentiation of male BJFF.6 or female AN1.1 iPSCs both with WT *TP53* induced the expression of MYOD1 and formed tumors in immunocompromised mice that histologically recapitulated solid-type ARMS (Supplemental Figure 2a-d). Since there was no change in the ability to form solid-type ARMS, we moved forward with WT *TP53* BJFF.6 cells for subsequent analyses.

Because P3F is a transcription factor and is thought to establish the overall expression signatures composing ARMS, we sought to determine its early, direct target genes upon P3F induction. For this task, we returned to our published iARMS system, minimally modified to use *TP53* wild-type BJFF.6 cells^17, 31^. We collected cells for analysis on Days 0 and 5 (prior to P3F transduction) and both P3F-transduced and un-transduced (UT) cells for analysis on Days 7, 9, 11, and 15 during and following endothelial-directed differentiation to analyze P3F binding, enhancer landscape, and gene expression (Figure 3a). We profiled enhancer landscapes and P3F binding using CUT&RUN for H3K27ac and HA-tag (P3F) respectively. At Day 15 of directed differentiation, new enhancers were present in P3F-transduced iARMS cells that were not present in UT cells (Figure 3b-c). These enhancers showed specific enrichment for P3F binding (Figure 3b-c). We next determined the set of P3F bound enhancers at Day 15 and tracked their kinetics of formation through the time course. Following P3F transduction on Day 5, we observed a progressive increase in P3F binding and enhancer signal through Day 15 at these sites (Figure 3d-e and Supplemental Figure 3a-c). Importantly, at Day 7, two days post P3F transduction, we observed an increase in H3K27ac signal at these sites destined to become iARMS-specific enhancers, which suggests that the altered enhancer landscape and resulting developmental reprogramming were mediated quickly upon P3F expression (Figure 3e).

**Figure 3.**
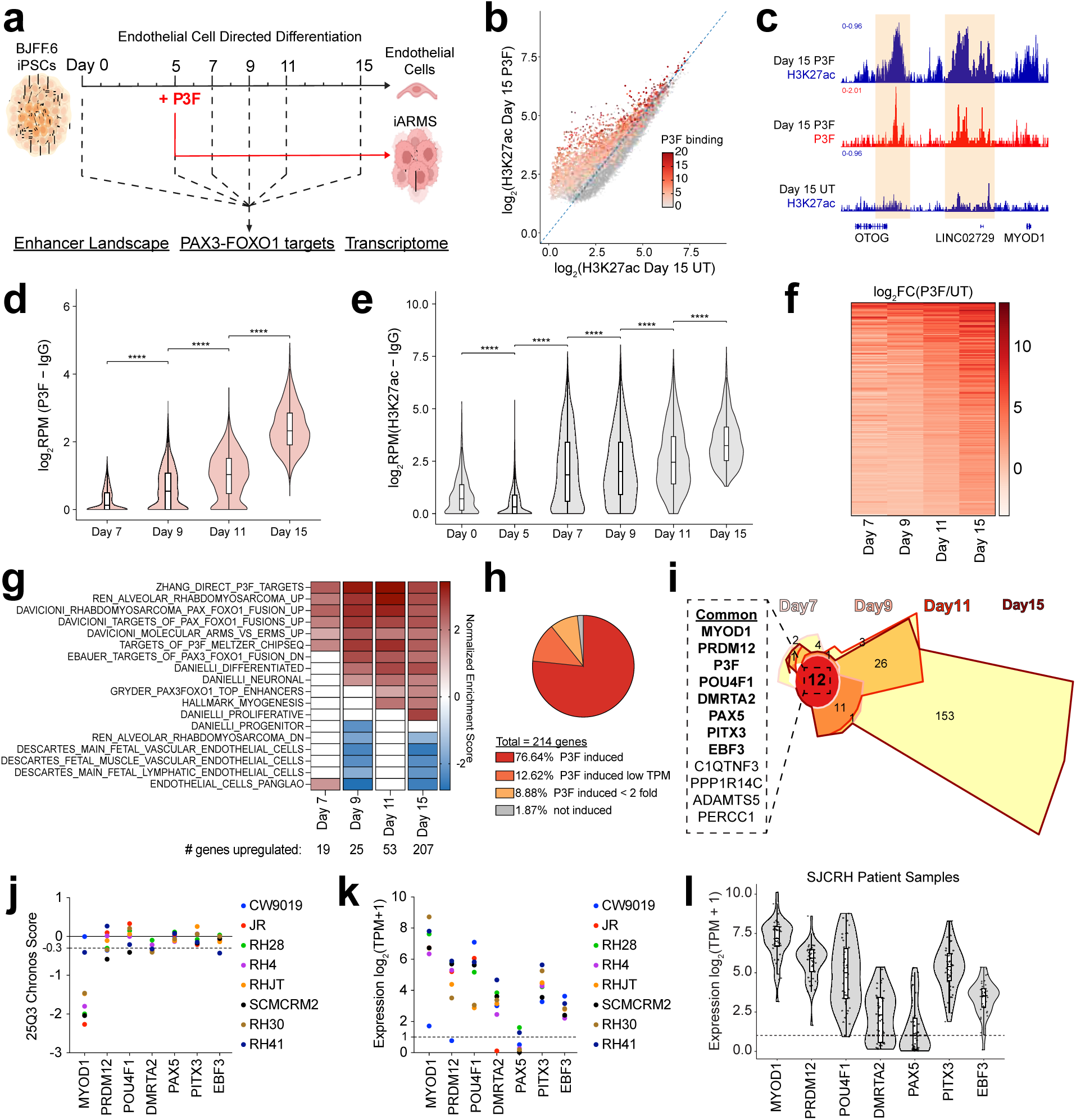
P3F establishes a novel transcriptional network to initiate tumorigenesis. **a)** Schematic of iARMS differentiation experiment with collection points labeled with dotted lines. **b)** Scatter plot of Day 15 H3K27ac signal in P3F-expressing cells (y-axis) and Day 15 H3K27ac signal in UT cells (x-axis). Signal was determined by log_2_(RPM). H3K27ac sites that are bound by P3F are colored in red scaled to the intensity of the P3F signal at each site. **c)** Representative traces at MYOD1 gene body and enhancers of Day 15 H3K27ac and Day 15 P3F CUT&RUN in iARMS and UT cells. **d)** Violin plot of P3F signal (log_2_RPM(P3F – IgG)) in each day of iARMS time course post transduction at genomic regions destined to become Day 15 P3F bound enhancers. **e)** Violin plot of H3K27ac signal (log_2_RPM(H3K27ac – IgG)) in each day of iARMS time course at genomic regions destined to become Day 15 P3F bound enhancers. **f)** Heatmap of significantly P3F-upregulated genes at Day 7, Day 9, Day 11, and Day 15 of the time course. Gene list was determined by combining significantly upregulated genes at Days 7, 9, 11, and 15. Each row is a gene, and the color represents the log_2_FC(P3F/UT) value at the corresponding time point. Genes are ranked by highest row mean to lowest row mean. **g)** Heatmap of Normalized Enrichment Scores (NES) of multiple gene sets for gene expression from Day 7, Day 9, Day 11, and Day 15 of the time course. Gene Set Enrichment Analysis was used to calculate the NES for each gene set for each sample pair (P3F/UT at corresponding time point). **h)** Pie chart displaying percentage of P3F upregulated genes that were categorized as either “P3F induced,” “P3F induced low TPM,” “P3F induced < 2 fold,” or “not induced.” **i)** Chow-Ruskey plot showing the common upregulated genes from Day 7, Day 9, Day 11, and Day 15 of the time course. Each edge is a time point sample and colored accordingly. The area of each segment is proportional to the number of genes in the depicted region. **j)** 25Q3 Chronos dependency scores from DepMap for iARMS time course early-response genes in ARMS cell lines. Each cell line is individually colored. **k)** Gene expression values for iARMS time course early-response TFs in ARMS cell lines from DepMap. Gene expression values are represented as log_2_(TPM+1) where expression is normalized as transcripts per million. **l)** Gene expression values for iARMS time course early-response TFs in P3F harboring ARMS patient tumors at diagnosis from St. Jude Cloud^43^.

### P3F rapidly upregulates select muscle and neural developmental TFs during iARMS initiation

To explore the consequences of P3F binding and enhancer activation on gene expression, we performed RNA-seq at Days 7, 9, 11, and 15 and compared the gene expression of the P3F cells to the UT cells at the same time points. Analysis of transcriptomes across the time course revealed that genes activated by P3F increase in number and degree of upregulation over the time course (Figure 3f and Supplemental Figures 3d-g). Two days following P3F transduction, only 19 genes were significantly upregulated at Day 7 compared to UT cells, but the overall expression levels of RMS gene sets were elevated, including that of ZHANG_DIRECT_P3F_TARGETS and REN_ALVEOLAR_RHABDOMYOSARCOMA_UP (Figure 3g and Supplemental Figure 3d)^14, 38^. Among the few early upregulated genes, the RMS diagnostic marker MYOD1 was the only fate-defining myogenic regulatory TF upregulated at Day 7 (Supplemental Figure 3d). Analyzing RNA-seq across our differentiation time course, we found an increased number of genes upregulated between P3F and UT cells at each time point and concomitantly an increase in the number of RMS gene sets significantly enriched (Figure 3f-g and Supplemental Figure 3d-g). Comparing our bulk transcriptome data to a recently published RMS cell identity atlas from single-cell RNA-seq data revealed that the “differentiated” (DANIELLI_DIFFERENTIATED) and “neuronal” (DANIELLI_NEURONAL) gene sets were the first RMS atlas gene sets to be significantly upregulated by P3F at Day 9, which is after we observed elevated expression of other established RMS signature gene sets. This upregulation was then followed by upregulation of the “proliferative” (DANIELLI_PROLIFERATIVE) gene set at Day 15 (Figure 3g)^35^. We never observed significant P3F-induced upregulation of the “progenitor” (DANIELLI_PROGENITOR) gene set when compared to our UT cells^35^. We sought to understand if the upregulated genes were truly induced after P3F expression or rather if their expression was maintained while decreasing during normal endothelial differentiation. Linear regression analysis of gene expression changes from Days 7, 9, 11, and 15 of differentiation revealed that only 4 genes were held consistent in P3F transduced cells while decreasing in untransduced cells (not induced, Figure 3h). All other relatively upregulated genes were among induced categories, indicating P3F expression indeed upregulated gene expression (P3F induced, P3F induced low TPM, and P3F induced < 2 fold, Figure 3h and Supplemental Figure 3h-k). These data suggest P3F induces specific transcriptional changes that progressively establish characteristics of an RMS cell identity expression program.

In addition to expression signatures of cellular identity, a putative core-regulatory circuit (CRC) was previously identified for ARMS cells^39^. We sought to understand the emergence of this circuit during the P3F-mediated transformation event. Analyzing RNA-seq across the differentiation time course, we observed P3F-induction of CRC members MYOD1, PITX3, and MYOG by Day 15, with only MYOD1 and PITX3 showing appreciable P3F-induced upregulation early in the time course (Supplemental Figure 3l-m). We were surprised that most putative ARMS CRC members were not appreciably upregulated by P3F during the time course, even while some TFs were already expressed during endothelial-directed differentiation (Supplemental Figure 3l-m). Most striking was the lack of *SOX8* regulation, as SOX8 was previously determined to be a central CRC node alongside MYOD1 and MYOG (Supplemental Figure 3l-m)^39, 40^. Discrepancies between models may be explained by notable differences in the methodologies used to explore RMS expression programs and their establishment and maintenance.

We sought to explore the requirement of SOX8 in ARMS using an orthogonal field-standard genetically engineered mouse model (GEMM) of P3F-mediated transformation. To this end, we developed a novel GEMM allowing for conditional deletion of *Sox8* through insertion of loxP sites flanking exons 1 and 2 of *Sox8* (*Sox8^flox^*, Supplemental Figure 3n). We inter-crossed *Sox8^flox^* mice into the field-standard ARMS GEMM, which uses *Myf6-Cre* to drive expression of *P3F* and conditional loss of *Trp53*^41^. We confirmed the presence of 5’ and 3’ *Sox8* loxP sites in *<u>M</u>yf6-_Cre;<u>P</u>ax3_^P3Fm/P3Fm^_;Tr<u>p</u>53_^flox/flox^_;Rosa26_^td<u>T</u>omato/tdTomato^_;<u>S</u>ox8_^+/+^* _(MPPTS_^WT^_), *Myf6-Cre;Pax3*_*^P3Fm/P3Fm^_;Trp53_^flox/flox^_;Rosa26_^tdTomato/tdTomato^_;Sox8_^flox/+^* _(MPPTS_^cHet^_), and *Myf6-Cre;Pax3*_*^P3Fm/P3Fm^_;Trp53_^flox/flox^_;Rosa26_^tdTomato/tdTomato^_;Sox8_^flox/flox^* _(MPPTS_^cKO^_) through PCR_ of isolated genomic DNA (Supplemental Figure 3o-p). We observed no significant difference between MPPTS^WT^, MPPTS^cHet^, and MPPTS^cKO^ tumor penetrance and a significant reduction in MPPTS^cKO^ tumor latency, suggesting that SOX8 is dispensable for RMS transformation (Supplemental Figure 3q). We confirmed recombination of the *Sox8^flox^*alleles through PCR of isolated tumor DNA and that expression of *Sox8* was significantly reduced in MPPTS^cKO^ tumors compared to MPPTS^WT^, both indicating successful *Sox8* deletion upon Cre-mediated recombination (Supplemental Figure 3r-s). Finally, tumors from MPPTS^WT^, MPPTS^cHet^, and MPPTS^cKO^ mice all displayed characteristic histologic features of solid variant ARMS with positive reactivity to MYOD1, MYOG, and DESMIN (Supplemental Figure 3t). These data suggest SOX8 is not required for initiating ARMS tumor cell identity and show that the iARMS model provides a unique platform to interrogate the essential TFs constructing ARMS cell identity.

We next sought to use our iARMS system to identify the genes that may play a central role in P3F-mediated tumor initiation. To this end, we analyzed which genes were upregulated by P3F at Day 7 that remained upregulated through Day 15. Twelve genes were upregulated at all four timepoints, seven of which were transcription factors (*MYOD1*, *PRDM12*, *POU4F1*, *DMRTA2*, *PAX5*, *PITX3*, and *EBF3*) (Figure 3i, TFs bolded). *MYOD1* was the only fate-defining myogenic regulatory TF consistently upregulated. All other TFs have been implicated in neural pathways, consistent with recent studies showing initial upregulation by P3F of neural genes in zebrafish embryos^42^. These genes do not collectively represent genetic dependencies in ARMS cell lines but are expressed in ARMS cell lines (Figure 3j-k). Furthermore, analysis of P3F-harboring ARMS tumors from St. Jude Children’s Research Hospital showed expression of these TFs in patient tumors at diagnosis (Figure 3l)^43^. These data show that a combination of myogenic and neural early response TFs is expressed during RMS initiation and are representative of genes expressed in human ARMS cell lines and patient tumors. This further demonstrates the capability of our iARMS system to recapitulate human ARMS and its utility to interrogate ARMS construction. However, it remains unclear if both muscle and neural TFs participate in ARMS tumor initiation or whether these TFs are simply a consequence of P3F transcription. Additionally, whether these TFs mark individual cell states or work in concert in the same cell cannot be appreciated from these bulk transcriptome data.

### Myogenic and neural TF expression is acquired rapidly after P3F expression

To better understand the contributions of myogenic and neural TFs to P3F-mediated transformation, we increased the resolution of our analyses by performing 10X single-cell Multiome (gene expression + ATAC-seq) profiling at the same time points during directed differentiation. As expected from our experimental design, not every cell from a given time point expressed P3F. wnnUMAP projection and clustering of all samples (n = 74,179 cells) revealed a diversity of clusters (Figure 4a). Cells from most samples inhabited multiple clusters, yet P3F expression was detected mainly in clusters 6 and 14 (Figure 4b and Supplemental Figure 4a-b). Accordingly, single-sample GSEA revealed that cells in clusters 6 and 14 showed elevated collective expression of RMS gene sets, including REN_ALVEOLAR_RHABDOMYOSARCOMA_UP and ZHANG_DIRECT_TARGETS (Figure 4c-d). Importantly, clusters 6 did not show expression of characteristic endothelial gene sets (Figure 4e). Given virtually all P3F-expressing cells were in clusters 6 and 14 showing low inter-cluster or intra-cluster heterogeneity, we concluded that clusters 6 and 14 represent RMS cells, and that there is no observable intermediate stage between endothelial precursor and P3F-expressing RMS-like cell beginning two days post-P3F expression.

**Figure 4.**
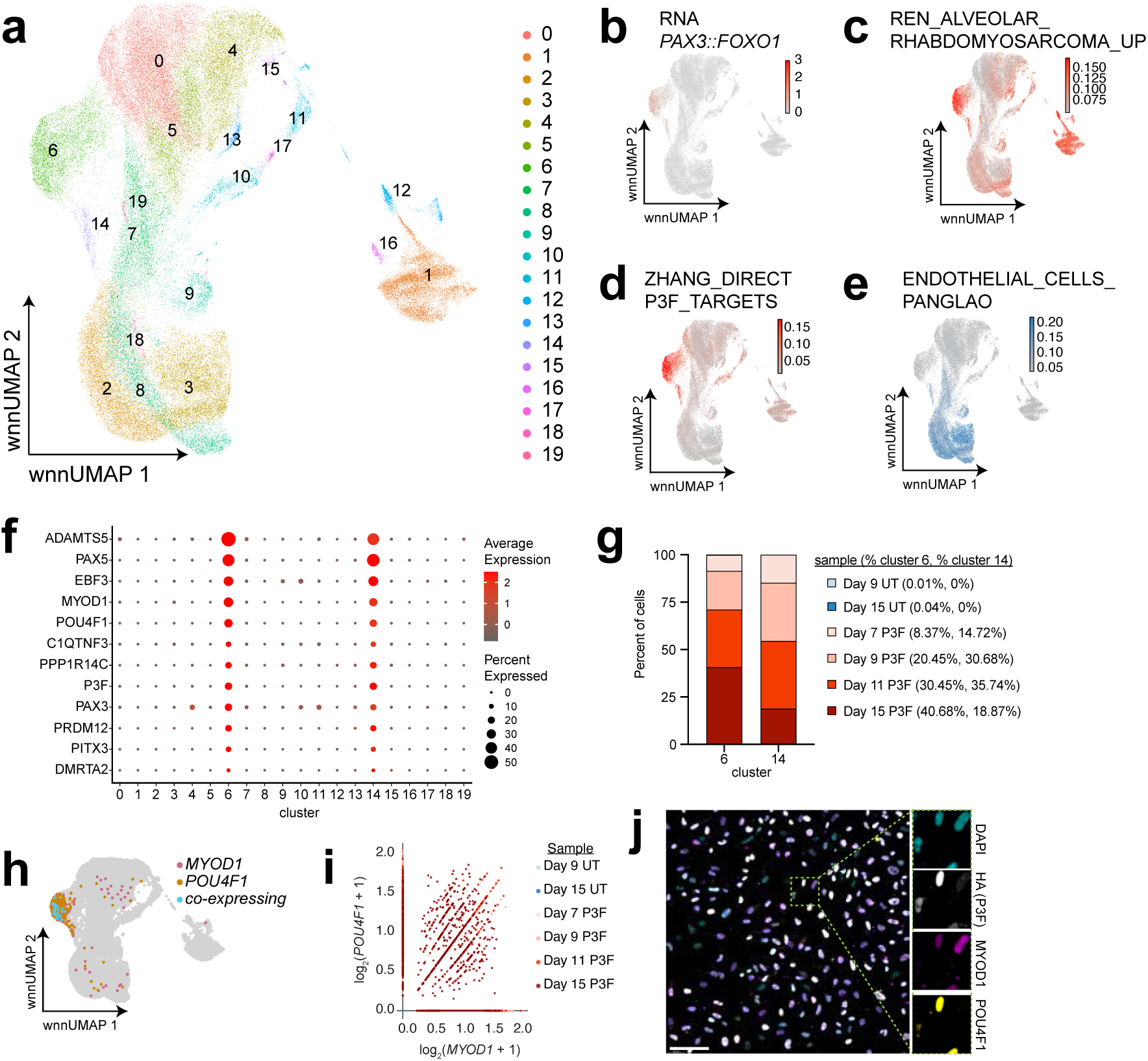
ARMS cell identity is acquired quickly through expression of both myogenic and neural TFs. **a)** wnnUMAP projection and cluster analysis of single-cell multiome (gene expression + ATAC-seq) profiles of Day 0, Day 5, Day 7 UT and P3F, Day 9 UT and P3F, Day 11 UT and P3F, and Day 15 UT and P3F samples. **b)** wnnUMAP highlighting *P3F* expression. **c)** wnnUMAP highlighting cells enriched for RMS gene set REN_ALVEOLAR_RHABDOMYOSARCOMA_UP. Enrichment score determined by single-sample GSEA. **d)** wnnUMAP highlighting cells enriched for RMS gene set ZHANG_DIRECT_P3F_TARGETS. Enrichment score determined by single-sample GSEA. **e)** wnnUMAP highlighting cells enriched for Endothelial gene set downloaded from PanglaoDB. Enrichment score determined by single-sample GSEA. **f)** Dot plot of gene expression in all clusters. Gene list defined as genes commonly upregulated in bulk transcriptomic data (Figure 3G). **g)** Stacked bar graph showing the proportion of cells from each sample for clusters 6 and 14. **h)** wnnUMAP highlighting cells expressing exclusively *MYOD1* (pink), exclusively *POU4F1* (orange), or both (cyan). **i)** Scatter plot showing expression of *MYOD1* (x-axis) and *POU4F1* (y-axis) in each cell from cluster 6. Each dot is an individual cell. Each cell is colored by its time course sample. **j)** Representative immunofluorescence of iARMS cells for DAPI (cyan), P3F (gray), MYOD1 (magenta), and POU4F1 (yellow). Scale bar = 100 µm.

To refine our iARMS cell populations, we compared our data to the recently published RMS identity single-cell atlas^35^. Cells in clusters 6 and 14 showed elevated expression of the DANIELLI_DIFFERENTIATED gene set, but intriguingly this gene set also showed expression in cells from cluster 1 that are solely from Day 0 of differentiation when the cells were still pluripotent and did not express P3F (Supplemental Figure 4c). The DANIELLI_DIFFERENTIATED gene set was the only RMS atlas gene set that correlated strongly with cells that expressed P3F and known RMS signatures in our data. The other three RMS atlas gene sets more accurately reflected clusters and cells that did not express P3F (Supplemental Figure 4c-f). Therefore, we turned our analyses to specifically explore P3F-expressing clusters 6 and 14.

The twelve early-response P3F-upregulated genes identified in our bulk expression analyses were uniquely expressed in clusters 6 and 14 (Figure 4f). Clusters 6 and 14 similarly express the early-response genes, indicating that the myogenic and neural TFs were not responsible for two independent cell states but were instead fundamental to both cell populations. Targeted analysis of clusters 6 and 14 revealed these clusters consist of P3F-expressing cells from each P3F-transduced timepoint (Figure 4g). Furthermore, we examined the sequences under ATAC-seq peaks from cells in each cluster and found that the binding motif for P3F was enriched in ATAC-seq peaks from cells in clusters that expressed P3F (Supplemental Figure 4g-h). Additionally, we observed the binding motif for MYOD1 in ATAC-seq peaks from cells that express P3F, supporting our notion that P3F induces a novel identity quickly with the aid of TFs like MYOD1 (Supplemental Figure i-j).

Since clusters 6 and 14 largely contained cells that expressed myogenic and neural TFs of interest, we next sought to determine if these TFs were expressed in the same cells. We plotted expression of the top myogenic (*MYOD1*) and neural (*PRDM12* and *POU4F1*) TFs in single cells from clusters 6 and 14 and discovered that myogenic and neural TFs were frequently co-expressed, especially in cluster 6 (Figure 4h-i and Supplemental Figure 4k-n). Immunofluorescence staining of iARMS cells for MYOD1 and POU4F1 similarly revealed individual cells that express only MYOD1, only POU4F1, or both TFs (Figure 4j and Supplemental Figure 4o). Taken together, these data show that RMS identity is quickly acquired upon expression of P3F and that this new identity is accompanied with the expression and activity of both myogenic and neural TFs.

### MYOD1 and POU4F1 are required for iARMS identity and not required for P3F-mediated transformation

To test whether these early-response TFs were required for transformation and/or establishment of iARMS cell identity, we created knockout (KO) BJFF.6 iPSC clones for each of *POU4F1*, *PRDM12*, and *MYOD1* (collectively referred to as TF-KO iPSCs). Before putting the TF-KO cells through our iARMS differentiation protocol, we confirmed the TF-KO iPSCs retained pluripotency through the ability to differentiate into all three germ layers. We subjected all TF-KO iPSCs to a germ layer differentiation protocol and determined that all TF-KO iPSCs were able to successfully differentiate into ectoderm (marked by OTX2 expression), mesoderm (marked by TBXT aka T expression), and endoderm (marked by SOX17 expression) (Supplemental Figure 5a-c).

We next subjected the TF-KO iPSCs to our standard iARMS differentiation protocol. Morphologic analysis of the cells after directed differentiation showed that all TF-KO iARMS cells resembled WT iARMS control cells (Figure 5a). All knockouts were successfully generated, as shown by expression of each gene being significantly reduced in the respective KO cell line with similar or higher expression of P3F compared to WT iARMS (Supplemental Figure 5d-g). To test the capacity for transformation, we performed a focus-formation assay with each TF-KO iARMS line. All lines successfully formed foci indicating TF-KO iARMS were able to transform (Figure 5b).

**Figure 5.**
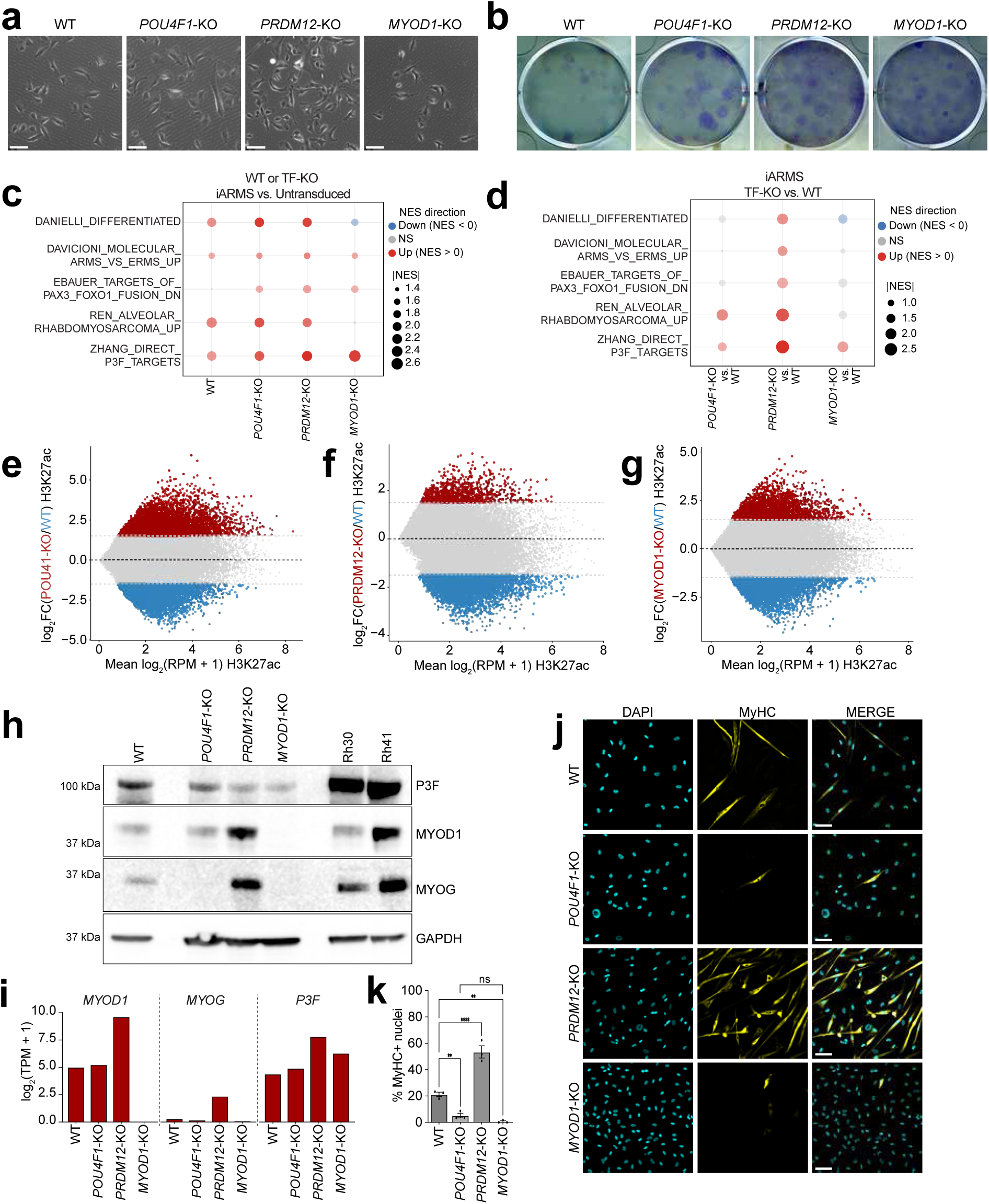
MYOD1 and POU4F1 are required for RMS identity and not required for P3F-mediated transformation. **a)** Phase contrast images of TF-KO and WT iARMS cells in culture. Scale bar = 100 µm. **b)** Representative wells of a focus-formation assay showing foci (purple) in WT iARMS and TF-KO iARMS cell lines. n = 3 wells **c)** Dot plot showing significant NES for multiple ARMS gene sets for each Day 15 TF-KO iARMS sample (WT, *POU4F1*-KO, *PRDM12*-KO, and *MYOD1*-KO) compared to the respective UT sample. Size of dots determined by absolute NES score. Opacity of dot determined by adjusted p-value if significant (adjusted p-value <0.05). Color of dot represents direction of NES (red = up, blue = down). **d)** Dot plot showing significant NES for multiple ARMS gene sets for each Day 15 TF-KO iARMS sample (*POU4F1*-KO, *PRDM12*-KO, and *MYOD1*-KO) compared to Day 15 WT iARMS. Size of dots determined by absolute NES score. Opacity of dot determined by adjusted p-value if significant (adjusted p-value <0.05). Color of dot represents direction of NES (red = up, blue = down). **e)** MA plot of mean log_2_(RPM + 1) H3K27ac signal (x-axis) versus log_2_FC(*POU4F1*-KO/WT) H3K27ac signal (y-axis) showing enhancers enriched for either sample (WT = blue, *POU4F1*-KO = red). **f)** MA plot of mean log_2_(RPM + 1) H3K27ac signal (x-axis) versus log_2_FC(*PRDM12*-KO/WT) H3K27ac signal (y-axis) showing enhancers enriched for either sample (WT = blue, *PRDM12*-KO = red). **g)** MA plot of mean log_2_(RPM + 1) H3K27ac signal (x-axis) versus log_2_FC(*MYOD1*-KO/WT) H3K27ac signal (y-axis) showing enhancers enriched for either sample (WT = blue, *MYOD1*-KO = red). **h)** Immunoblot of WT iARMS, *POU4F1*-KO iARMS, *PRDM12*-KO iARMS, *MYOD1*-KO iARMS, Rh30, and Rh41 cells for P3F, MYOD1, MYOG, and GAPDH. **i)** Gene expression values from RNA sequencing for *MYOD1*, *MYOG*, and *P3F* for WT, *POU4F1*-KO, *PRDM12*-KO, and *MYOD1*-KO iARMS. Gene expression is represented as log_2_(TPM +1). **j)** Representative immunofluorescence of WT iARMS, *POU4F1*-KO iARMS, *PRDM12*-KO iARMS, and *MYOD1*-KO iARMS for DAPI (cyan) and MyHC (yellow) after culturing in myogenic differentiation medium for 5 days. n = 3 biological replicates. **k)** Differentiation index calculation of WT, *POU4F1*-KO, *PRDM12*-KO, and *MYOD1*-KO iARMS cultured in myogenic differentiation medium for 5 days. n = 3 biological replicates. All p-values in pairwise comparisons were determined by Student’s t-test (unpaired, two-tailed); *p < 0.05, **p < 0.01, ***p < 0.001, ****p < 0.0001. Data represented as mean +/- SEM.

To explore if the TF-KO iARMS had any transcriptional changes compared to WT iARMS, we performed RNA sequencing at Day 15 of iARMS directed differentiation. In P3F-transduced samples compared to their respective UT control, we observed a significant but inconsistent upregulation of ARMS gene sets by P3F in both WT and TF-KO iARMS cells (Figure 5c). Interestingly, *MYOD1*-KO iARMS cells were the only condition to show a lack of P3F-induced upregulation of the DANIELLI_DIFFERENTIATED gene set and REN_ALVEOLAR_RHABDOMYOSARCOMA_UP gene set, indicating a potential failure to acquire ARMS identity without MYOD1 despite the presence of P3F (Figure 5c and Supplemental Figure 5d). Comparing each TF-KO iARMS cell line to WT iARMS revealed findings consistent in regard to *MYOD1*-KO iARMS (Figure 5d). Interestingly, *POU4F1*-KO iARMS cells only showed upregulation of two gene sets when compared to WT iARMS, indicating *POU4F1*-KO iARMS may have altered iARMS identity acquisition similar to *MYOD1*-KO iARMS cells (Figure 5d). Surprisingly, *PRDM12*-KO iARMS cells showed upregulation of all five gene sets compared to WT iARMS cells, indicating that PRDM12 may function as a repressor of iARMS identity (Figure 5d). We next sought to understand if the gene expression changes were reflective of different enhancer landscapes. We performed CUT&RUN for H3K27ac in each TF-KO iARMS cell line and compared the enhancers to WT iARMS at Day 15 of directed differentiation. Consistent with changes in gene expression, each TF-KO iARMS cell line had altered enhancer landscapes compared to WT iARMS (Figure 5e-g). One notable region contains increased H3K27ac signal in TF-KO iARMS surrounding *MYF5* and *MYF6* when compared to WT iARMS (Supplemental Figure 5h). This response could indicate a potential compensation mechanism, and thus we sought to investigate markers of ARMS at the protein level.

Immunoblotting of WT and TF-KO iARMS cells revealed that both *POU4F1*-KO and *MYOD1*-KO iARMS cells do not express the RMS diagnostic marker MYOG, confirming compromised iARMS identity and suggesting impaired myogenic differentiation ability in their absence (Figure 5h). Interestingly, *POU4F1*-KO iARMS cells showed no MYOG expression while maintaining expression of MYOD1. Conversely, *PRDM12*-KO iARMS cells showed increased expression of MYOD1 and MYOG when compared to WT iARMS cells (Figure 5h). These trends were also seen at the level of gene expression (Figure 5i). Importantly, these expression changes reflected the differentiation potential of TF-KO iARMS cells and altered ARMS identity. When cultured in muscle differentiation media for 5 days, *POU4F1*-KO and *MYOD1*-KO iARMS cells showed a significant reduction in myogenic differentiation potential, while *PRDM12*-KO iARMS cells showed a significant increase in myogenic differentiation potential compared to WT iARMS (Figure 5j-k). Interestingly, xenografted TF-KO iARMS cells all resulted in solid-type ARMS tumors in immunocompromised mice (Supplemental Figure 5i). These data could indicate that P3F can induce genetic compensation of alternative myogenic and/or neural factors to aid in ARMS establishment. During muscle development MYF5 and MYOD1 play redundant roles and have been previously shown to compensate for the absence of each other^44^. Indeed, gene expression analysis by qPCR and immunoblotting confirmed increased expression of MYF5 in TF-KO iARMS tumors compared to WT iARMS tumors (Supplemental Figure 5j-k). These data could also suggest that additional secreted factors or supporting cells absent in cell culture systems will be important for future dissection of ARMS establishment. Taken together, these data indicate that both POU4F1 and MYOD1 are required for establishment of iARMS cell identity, while PRDM12 may act as a differentiation repressor. Additionally, our data suggest that neural TFs may act in modulating the myogenic state of iARMS cells. Furthermore, our data show that P3F can act upon other myogenic regulatory factors in the absence of these important TFs. In total, our data reveals POU4F1 and MYOD1 to be essential TFs for acquisition of iARMS identity and PRDM12 as an important differentiation repressor in iARMS cells.

## Discussion

In this study, we sought to explore the separate roles of P3F in RMS initiation and identity through the generation and integration of multiple human stem cell models of alveolar rhabdomyosarcoma (ARMS). We generated a novel human induced pluripotent stem cell-derived model of ARMS by expressing a doxycycline-inducible, degradable P3F-FKBP12^F36V^ during endothelial-directed differentiation (ddP3F cells). This model allowed for fine control over P3F expression during and after transformation. This new ddP3F system complements our published iARMS model by incorporating ectopic expression of P3F with an epitope tag that provided unique opportunities to specifically identify P3F-expressing cells and a robust means to measure P3F binding. Our data show that “P3F off” ddP3F cells were still viable, continued to proliferate, and retained the ability to form tumor-like cell culture foci, though they had reduced myogenic differentiation capacity. Interestingly, while “P3F off” ddP3F cells did show changes in gene expression compared to “P3F on” ddP3F cells, our results show that ddP3F cells retained expression of gene programs associated with ARMS cell identity and could form alternative sarcomatous malignancies in vivo. We interpret that P3F is required for initiating but not maintaining ARMS cell identity.

We explored the mechanism of P3F-mediated ARMS initiation using our previously published iARMS system^31^. Our results demonstrated that P3F initiates a novel epigenetic program that results in the rapid and progressive emergence of ARMS cell identity. Further investigation revealed that upregulation of neural-fate-defining transcription factors was concomitant with ARMS identity acquisition. Surprisingly, very few members of the putative CRC proposed for ARMS were P3F-induced during the time course and instead may reflect normal cell-of-origin expression^39, 40^. Single-cell analysis during the transformation event revealed frequent co-expression of myogenic and neural transcription factors in individual cells. We ultimately concluded that the myogenic and neural transcription factors are individually regulating aspects of ARMS identity but not transformation.

Investigation of gene expression during the iARMS transformation time course revealed early and sustained upregulation by P3F of neural fate-defining TFs *POU4F1* and *PRDM12* in addition to muscle fate-defining TF *MYOD1*. *MYOD1* was the only muscle fate-defining TF induced by P3F in all time points, suggesting an importance in RMS identity-initiation. *MYOD1* deletion prevented the acquisition of the ARMS cell identity and showed functional impairment of myogenesis in vitro, with potential compensation from MYF5 upon xenografting. MYOD1 is the master regulator of muscle development, capable itself of transdifferentiating multiple cell types to muscle cells^45, 46^. Furthermore, MYOD1 has been shown to suppress cell death in RMS cells and play an important repressive role in fibroblast to muscle transdifferentiation^47^. Our data suggest that MYOD1 is an essential factor to establish RMS cell identity.

A current question in the RMS community centers around the source and relevance of neural expression signatures identified in human ARMS samples^48^. We identified six transcription factors that function in neuronal development that were among the twelve early-response genes across all time points. These TFs were not unique to our iARMS system as we observed expression in both human ARMS cell lines and patient tumors at diagnosis. We explored the function of the two most highly expressed neural TFs, POU4F1 and PRDM12, by creating KO iPSC lines, and observed opposing phenotypes with *POU4F1*-KO and *PRDM12*-KO cells. *POU4F1*-KO iARMS cells maintained MYOD1 expression but failed to express MYOG and complete myogenic differentiation. This suggests that POU4F1 and MYOD1 act cooperatively to initiate and maintain RMS cell identity. POU4F1 has previously been implicated in developing cardiomyocytes through regulation of Hsp27, but not in conjunction with MYOD1^49^. These results are unexpected, as MYOD1 is the master regulator of muscle differentiation. Additionally, our data reveal that PRDM12 may play an important repressive role in RMS. *PRDM12*-KO iARMS cells express more MYOD1 and MYOG resulting in an increased ability to undergo myogenic differentiation, suggesting that PRDM12 may act to normally restrain the full muscle-determining activity of MYOD1 in RMS cells, perhaps repressing overall differentiation rate and the maintenance of a suitably large fraction of undifferentiated cells. PRDM12 is expressed during neuronal development to establish nociceptive neurons but is largely not expressed in muscle or post-natal tissues^50, 51^. It has long been questioned why RMS cells fail to terminally differentiate while maintaining high expression of MYOD1 and MYOG. Our data have uncovered a potentially novel mode of MYOD1 regulation in RMS cells through cooperation with and repression by neural transcription factors that would otherwise not play a role in normal muscle development. In this way, P3F can borrow aspects of PAX3’s roles in normal muscle and normal neuronal development to establish a corrupt transcriptional network that initiates and maintains RMS tumors. Furthermore, we posit that the rapid establishment of both myogenic and neural networks specifically by P3F early response genes are essential for correct fate acquisition. Our findings suggest a potential mechanism of transcriptional control in ARMS that will require further validation in established models.

Previous studies have shed light on multiple cell states comprising individual RMS tumors and putative CRCs governing RMS identity as a whole^32–35, 39, 40^. These studies have been limited by focusing on deconstructive approaches like genetic dependencies and technical issues such as the inability to detect the P3F breakpoint using certain single-cell sequencing platforms. Nevertheless, these studies have been instrumental in our understanding of tumor heterogeneity and transcriptional networks in RMS, and thus, we sought to use our iARMS system to determine the order of operations necessary for the establishment of cell identity and transcriptional networks. While our iARMS system relies on ectopic expression of P3F with an epitope tag, this potential limitation prov ided a constructivist opportunity to specifically identify the P3F-expressing cells at single-cell resolution and a robust means to measure P3F genome occupancy. We found that P3F establishes novel enhancers resulting in expression of myogenic and neural genes that do not define distinct cell states but rather reflected the presence of P3F and its collaborating TFs at both bulk and single-cell resolutions. Comparing our data to a recently defined RMS cell atlas, we observed that only the “differentiated” cell state accurately described P3F-expressing cells in our system. As expected, P3F direct target gene expression was only detected in the P3F-expressing, “differentiated” cell state. The “proliferative,” “progenitor,” and “neural” cell states were all enriched in populations not expressing P3F. Interestingly, these states were enriched in the pluripotent stem cells from Day 0 prior to endothelial-directed differentiation. This could signify the possibility of stem cell like features of RMS cells. Furthermore, when we looked for establishment of previously defined putative CRC members, the only consensus TF upregulated by P3F was MYOD1. Additional exploration into putative core CRC hub TF SOX8 revealed no impact on RMS cell identity or tumor burden when deleted in an ARMS GEMM. Taken together, these data reveal the value of our iARMS system in refining the TFs responsible for establishing RMS cell identity including TFs not currently defined as genetic dependencies.

Our investigations of P3F-mediated initiation events were prompted by our observation that “P3F off” ddP3F cells continued to proliferate for over 20 passages, maintained focus-formation ability, and retained RMS gene expression programs upon loss of P3F. This finding is congruent with the observation from Pandey et al. that showed ARMS tumors in xenografts from genetically engineered human myoblasts recurred without expressing P3F^28^. These data demonstrate the stability of the P3F-initiating tumorigenic gene expression program even after P3F is lost. With the advent of molecular glues and proteolysis targeting chimeras (PROTACs), therapeutically targeting oncogenic transcription factors has become tantalizing and feasible. Our data should not dissuade active investigation of P3F targeting molecular glues and PROTACs. However, exploration using model systems as presented herein will allow discovery of P3F-dependent and P3F-independent vulnerabilities, thus providing mechanistic insight into how children’s tumors could recur after targeting P3F as well as orthogonal targets that may enable synergistic targeting strategies.

## Limitations of the study

Our study relies on the constitutive expression of epitope-tagged P3F, and thus we cannot comment on the endogenous regulation or potential for P3F expression pulses. P3F expression in mouse tumors has been shown to change through cell cycle progression, which could be important for RMS cell state initiation^52^. Future studies with our ddP3F model with fine-tuned control of P3F could illuminate potential important facets of P3F expression pulses and expression levels on RMS cell state. Additionally, our multiomic profiling of iARMS initiation was only performed on cells in vitro. This limits our ability to determine if the identified early response genes co-express in xenografted tumor cells and the cell states therein. It will be interesting to incorporate single-cell multiomic data from xenografted iARMS tumor cells and patient derived xenografts into our current trajectory in future studies. Furthermore, one limitation of our single-cell multiomic assay is the difficulty in comprehensively capturing transcripts in a given cell or cells; therefore, we note that we may not have fully captured all P3F expressing cells in our dataset.

---

**Supplemental Figure 1.**
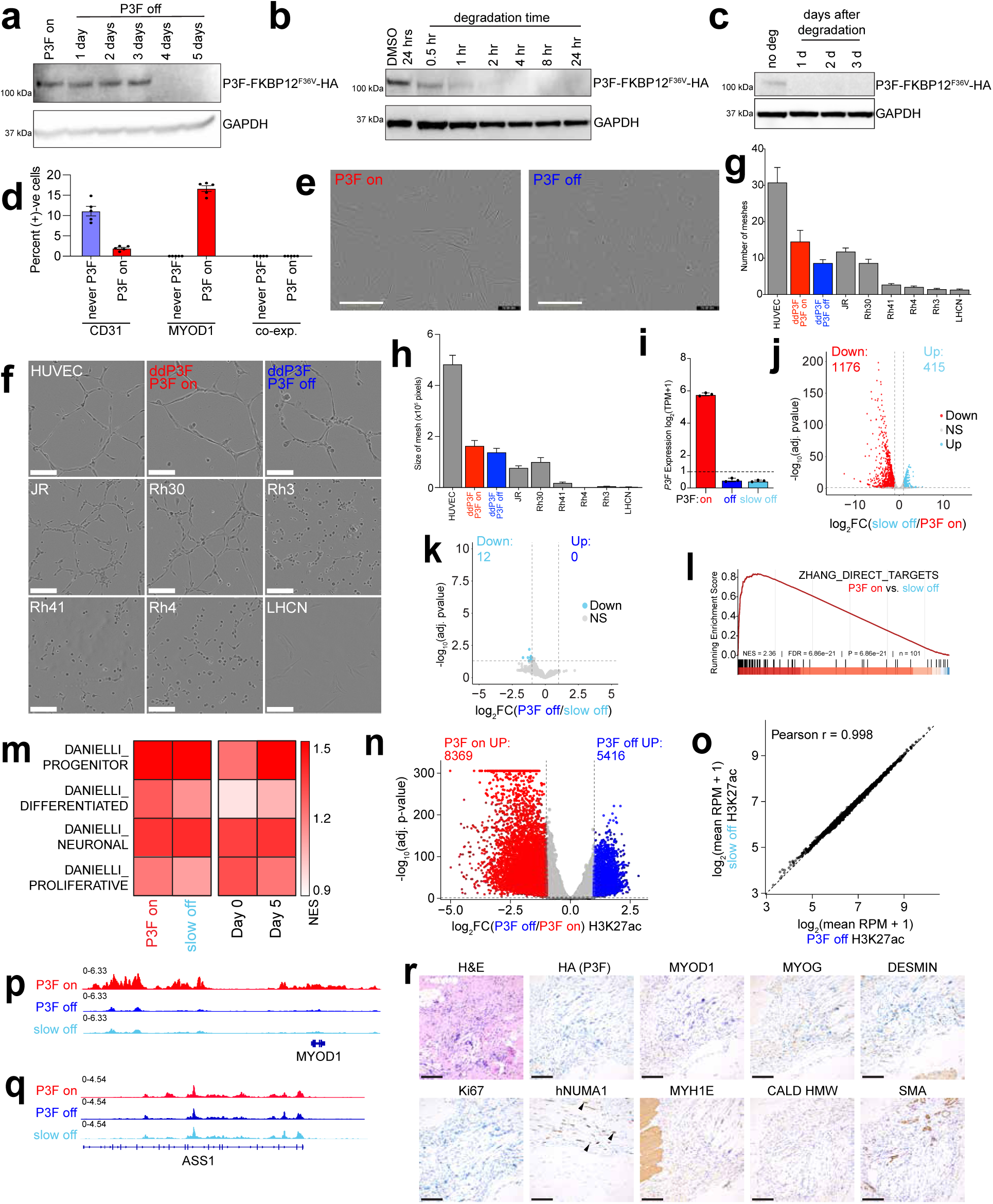
Functional validation of ddP3F “P3F on” and “P3F off” cells. **a)** Immunoblots of PAX3::FOXO1 and corresponding GAPDH in a time course following doxycycline removal of ddP3F cells cultured in doxycycline-free media. **b)** Immunoblots of PAX3::FOXO1 and corresponding GAPDH on a time course of doxycycline removal and addition of 100 µM dTAG^V^-1 in ddP3F cells. **c)** Immunoblots of PAX3::FOXO1 and corresponding GAPDH of ddP3F cells prior and a time course following doxycycline removal and 24 hours treatment with 100µM dTAG^V^-1 then removal and subsequent maintenance in doxycycline free media. Time reflects the number of days following dTAG^V^-1 removal. **d)** Quantification of CD31, MYOD1, and CD31/MYOD1 co-expressing cells in “never P3F” and “P3F on” ddP3F cells on Day 15 of endothelial directed differentiation. Data represented as mean +/- SEM. n = 5 images **e)** Representative Incucyte images from day 7 of proliferation measurements for “P3F on” and “P3F off” ddP3F cells. Scale bar = 400 µm. **f)** Representative images of various cell lines in an endothelial tube formation assay. Scale bar = 200 µm. **g)** Quantification of number of meshes in endothelial tube assay 8 hours after assay start (n = 3, 25 images analyzed for each cell line). Data represented as mean +/- SEM. **h)** Quantification of the size of meshes in endothelial tube assay 8 hours after assay start (n = 3 biological replicates, 25 images analyzed for each cell line). Data represented as mean +/- SEM. **i)** Gene expression by RNA sequencing of *P3F* in ddP3F “P3F on,” “P3F off,” and “slow off” cells. Gene expression was normalized as TPM and plotted as log_2_(TPM+1) and dashed line at y = 1 indicates expression threshold. **j)** Volcano plot showing changes in differentially expressed genes with absolute log_2_FC greater than 1 and false discovery rate (FDR) < 5% in ddP3F “P3F on” cells (red) and ddP3F cells cultured in doxycycline free media for 1 week (“slow off”, light blue). **k)** Volcano plot showing changes in differentially expressed genes with absolute log_2_FC greater than 1 and FDR < 5% in ddP3F “slow off” cells (light blue) and ddP3F “P3F off” cells at 1 week (blue). **l)** Enrichment plot of P3F-direct target gene set (ZHANG_DIRECT_P3F_TARGETS identified by Zhang et al.) for “slow off” ddP3F cells compared to “P3F on” ddP3F cells. **m)** Heatmap of the single sample gene expression of “slow off” ddP3F cells, “P3F on” ddP3F cells, and Day 0 and Day 5 of endothelial directed differentiation cells. Single-sample gene set enrichment analysis (ssGSEA) was used to score. ssGSEA was used to score proliferative (DANIELLI_PROLIFERATIVE), neuronal (DANIELLI_NEURONAL), differentiated (DANIELLI_DIFFERENTIATED), and progenitor (DANIELLI_PROGENITOR) cell states identified by Danielli et al. **n)** Volcano plot showing changes in H3K27ac signal with absolute log_2_FC greater than 1 and FDR < 5% in “P3F off” ddP3F cells (blue) and “P3F on” ddP3F cells (red). **o)** Scatter plot showing correlation of H3K27ac signal in “P3f off” ddP3F cells (x-axis) and “slow off” ddP3f cells (y-axis). H3K27ac signal is represented as log_2_(mean RPM + 1). **p)** Representative CUT&RUN alignment tracks for H3K27ac in “P3F on” ddP3F cells, “P3F off” ddP3F cells, and “slow off” ddP3F cells at the gene locus for *MYOD1* and its associated enhancers. **q)** Representative CUT&RUN alignment tracks for H3K27ac in “P3F on” ddP3F cells, “P3F off” ddP3F cells, and “slow off” ddP3F cells at the gene locus for *ASS1*. **r)** Representative histology (H&E) and immunohistochemistry (HA (P3F), MYOD1, MYOG, DESMIN, Ki67, hNUMA1, MYH1E, CALD HMW, and SMA) of “P3F stay off” ddP3F tumor bearing leg. Scale bar = 100 µm.

**Supplemental Figure 2.**
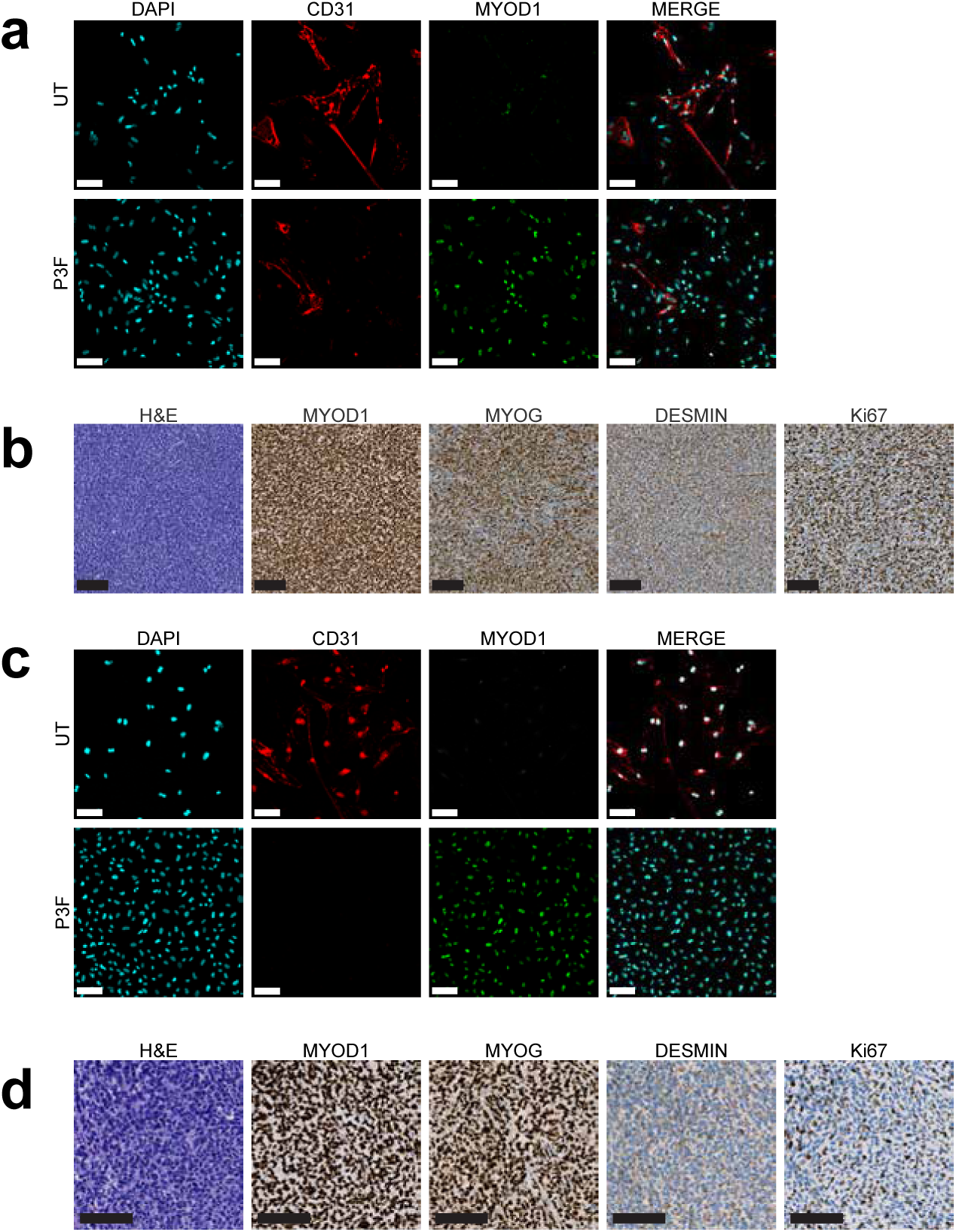
Generation of iARMS from *TP53* WT BJFF.6 male-derived iPSCs and AN1.1 female-derived iPSCs. **a)** Representative immunofluorescence of DAPI (cyan), CD31 (red), and MYOD1 (green) on cells at day 15 of endothelial-directed differentiation and iARMS cell generation from *TP53* WT BJFF.6 iPSCs. Scale bar = 100 µm. **b)** Representative histology (H&E) and immunohistochemistry (MYOD1, MYOG, DESMIN, and Ki67) of *TP53* WT BJFF.6 derived iARMS tumors. Scale bar = 100 µm. **c)** Representative immunofluorescence of DAPI (cyan), CD31 (red), and MYOD1 (green) on cells at day 15 of endothelial directed differentiation and iARMS cell generation from female-derived *TP53* WT AN1.1 iPSCs. Scale bar = 100 µm. **d)** Representative histology (H&E) and immunohistochemistry (MYOD1, MYOG, DESMIN, and Ki67) of female-origin *TP53* WT AN1.1 derived iARMS tumors. Scale bar = 100 µm.

**Supplemental Figure 3.**
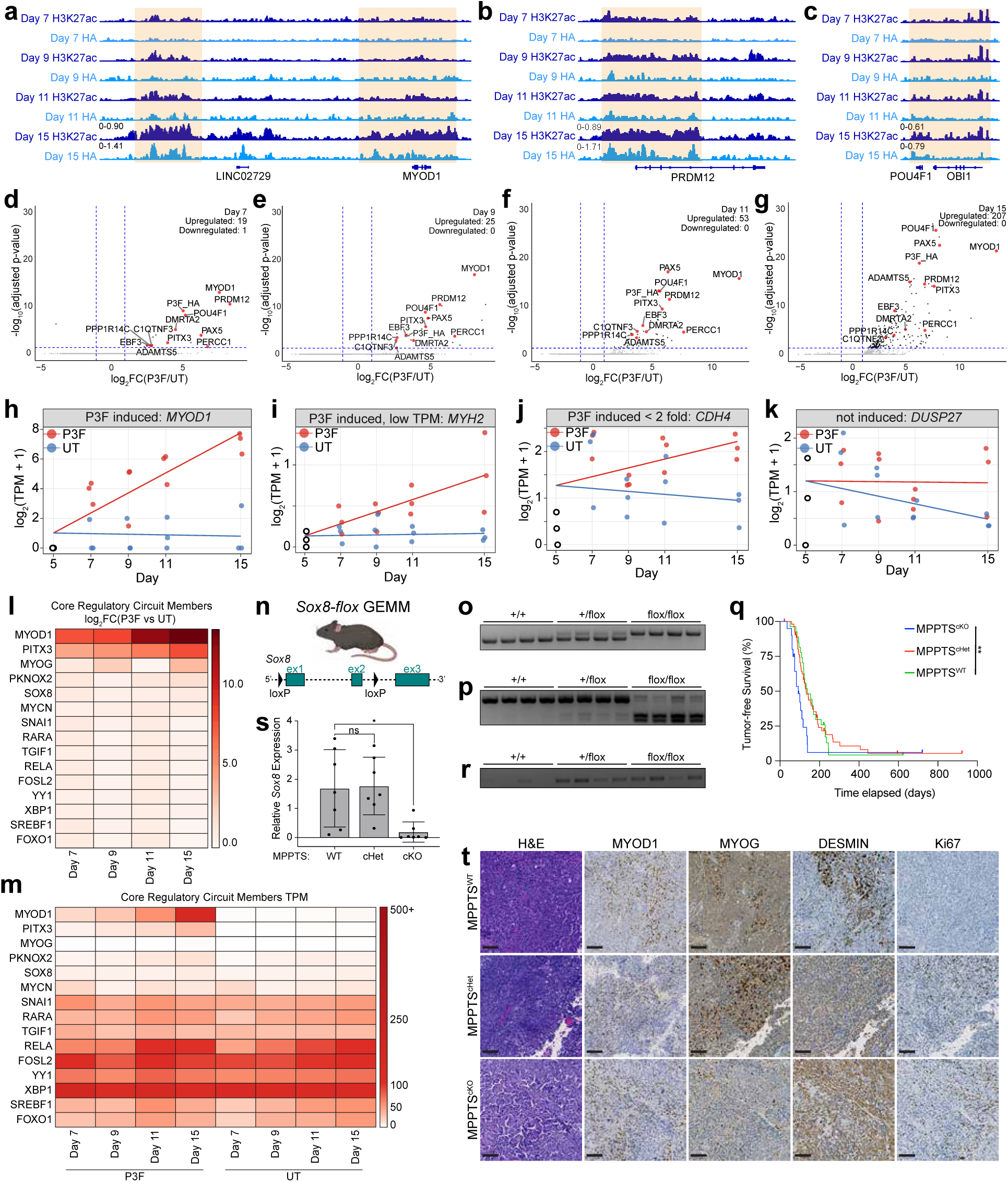
P3F establishes a novel transcriptional network to initiate tumorigenesis that does not contain previously identified core regulatory circuit members. **a)** CUT&RUN alignment tracks for H3K27ac and P3F in Day 7 P3F, Day 9 P3F, Day 11 P3F, and Day 15 P3F cells at the gene locus for MYOD1 and its associated enhancer. b) CUT&RUN alignment tracks for H3K27ac and P3F in Day 7 P3F, Day 9 P3F, Day 11 P3F, and Day 15 P3F cells at the gene locus for PRDM12. c) CUT&RUN alignment tracks for H3K27ac and P3F in Day 7 P3F, Day 9 P3F, Day 11 P3F, and Day 15 P3F cells at the gene locus for POU4F1. d) Volcano plot showing differentially expressed genes with absolute log_2_FC greater than 1 and false discovery rate (FDR) less than 5% in Day 7 P3F cells compared to Day 7 UT cells. Genes commonly upregulated in Day 7 P3F, Day 9 P3F, Day 11 P3F, and Day 15 P3F cells are colored in red. **e)** Volcano plot showing differentially expressed genes with absolute log_2_FC greater than 1 and false discovery rate (FDR) less than 5% in Day 9 P3F cells compared to Day 9 UT cells. Genes commonly upregulated in Day 7 P3F, Day 9 P3F, Day 11 P3F, and Day 15 P3F cells are colored in red. **f)** Volcano plot showing differentially expressed genes with absolute log_2_FC greater than 1 and false discovery rate (FDR) less than 5% in Day 11 P3F cells compared to Day 11 UT cells. Genes commonly upregulated in Day 7 P3F, Day 9 P3F, Day 11 P3F, and Day 15 P3F cells are colored in red. **g)** Volcano plot showing differentially expressed genes with absolute log_2_FC greater than 1 and false discovery rate (FDR) less than 5% in Day 15 P3F cells compared to Day 15 UT cells. Genes commonly upregulated in Day 7 P3F, Day 9 P3F, Day 11 P3F, and Day 15 P3F cells are colored in red. **h)** Representative linear regression scatter plot of “P3F induced” gene *MYOD1.* Criteria for “P3F induced” included TPM >1, log_2_FC > 2-fold in Day 15 P3F compared to UT, and overall positive slope in P3F cells compared to UT cells. **i)** Representative linear regression scatter plot of “P3F induced, low TPM” gene *MYH2.* Criteria for “P3F induced, low TPM” included TPM < 1, log_2_FC > 2-fold in Day 15 P3F compared to UT, and overall positive slope in P3F cells compared to UT cells. **j)** Representative linear regression scatter plot of “P3F induced < 2-fold” gene *CDH4.* Criteria for “P3F induced < 2-fold” included TPM > 1, log_2_FC < 2-fold in Day 15 P3F compared to UT, and overall positive slope in P3F cells compared to UT cells. **k)** Representative linear regression scatter plot of “not induced” gene *DUSP27.* Criteria for “not induced” included non-negative slope in P3F cells compared to negative slop in UT cells. **l)** Heatmap showing log_2_FC(P3F/UT) values of previously determined FP-RMS CRC TFs in Day 7 P3F, Day 9 P3F, Day 11 P3F, and Day 15 P3F samples. **m)** Heatmap showing TPM values of previously determined FP-RMS CRC TFs in Day 7, Day 9, Day 11, and Day 15 P3F and UT samples. **n)** Schematic of *Sox8-flox* GEMM showing insertion of loxP sites flanking exons 1 and 2 of *Sox8*. **o)** Genotyping gel showing presence of either WT or 5’ flox allele from representative mice. **p)** Genotyping gel showing presence of either WT or 3’ flox allele from representative mice. **q)** Kaplan-Meier tumor-free survival curve. MPPTS^WT^ (green, n = 20), MPPTS^cHet^ (red, n = 53), and MPPTS^cKO^ (blue, n = 20). Significance determined by Log rank Mantel-Cox text. **r)** Genotyping gel showing the presence of knock-out allele in representative mice. Non-specific band from WT mice was extracted and sequenced with the resulting sequence aligning to *Prag1*. **s)** Gene expression by real time PCR of *Sox8* in representative MPPTS^WT^, MPPTS^cHet^, and MPPTS^cKO^ tumors. **t)** Representative histology (H&E) and immunohistochemistry (MYOD1, MYOG, DESMIN, and Ki67) of MPPTS^WT^, MPPTS^cHet^, and MPPTS^cKO^ tumors. Scale bar = 100 µm. All p-values in pairwise comparisons were determined by Student’s t-test (unpaired, two-tailed); *p < 0.05. Data represented as mean +/- SEM.

**Supplemental Figure 4.**
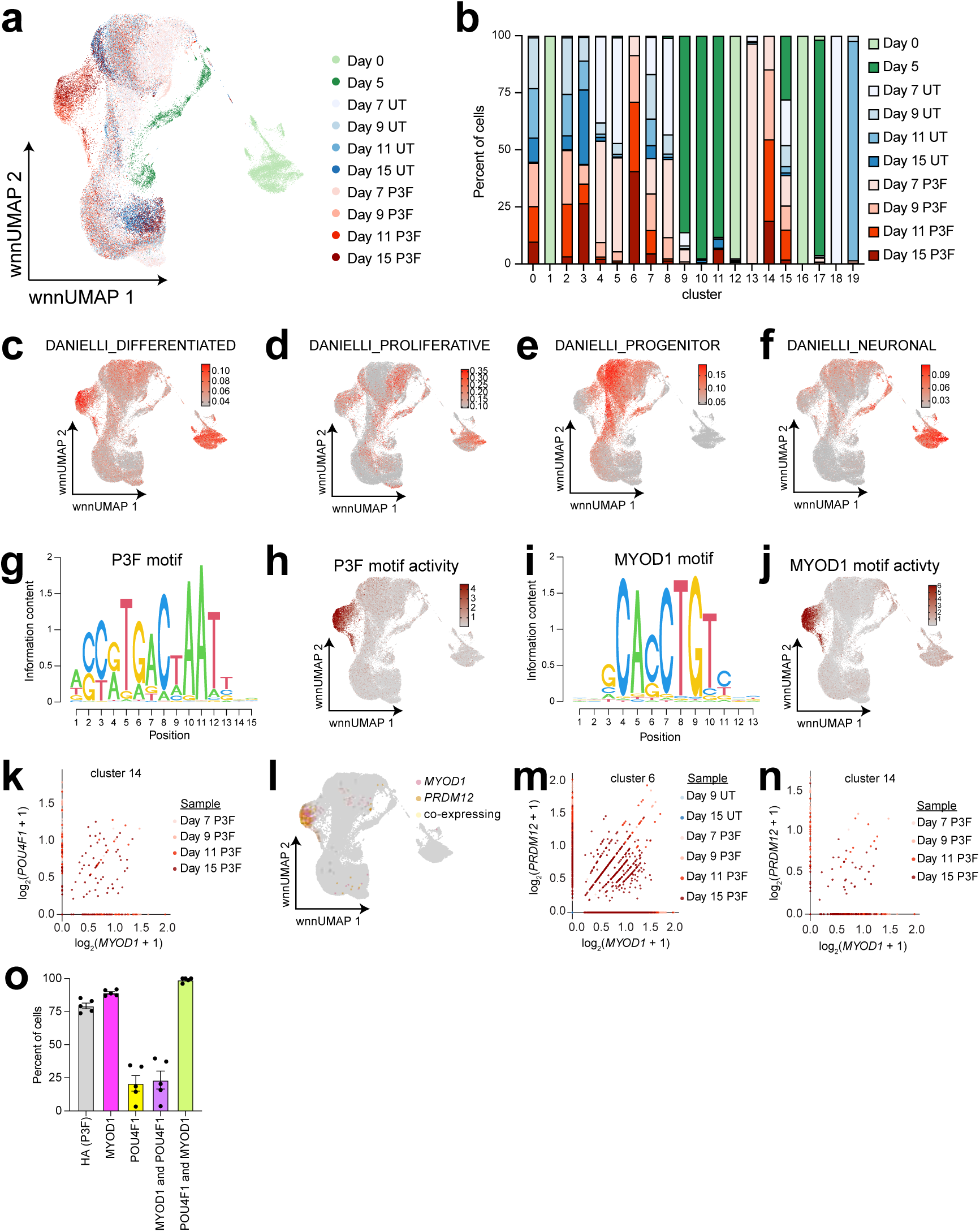
ARMS cell fate is acquired quickly through expression of both myogenic and neural TFs. **a)** wnnUMAP colored by sample. **b)** Stacked bar graph showing the proportion of cells from each sample for all clusters. **c)** wnnUMAP highlighting cells enriched for RMS atlas gene set DANIELLI_DIFFERENTIATED. Enrichment score determined by single-sample GSEA. **d)** wnnUMAP highlighting cells enriched for RMS atlas gene set DANIELLI_PROLIFERATIVE. Enrichment score determined by single-sample GSEA. **e)** wnnUMAP highlighting cells enriched for RMS atlas gene set DANIELLI_PROGENITOR. Enrichment score determined by single-sample GSEA. **f)** wnnUMAP highlighting cells enriched for RMS atlas gene set DANIELLI_NEURONAL. Enrichment score determined by single-sample GSEA. **g)** Position weight matrix depicting P3F binding motif. **h)** wnnUMAP highlighting cells with P3F binding motif activated in open chromatin. **i)** Position weight matrix depicting MYOD1 binding motif. **j)** wnnUMAP highlighting cells with MYOD1 binding motif activated in open chromatin. **k)** Scatter plot showing expression of *MYOD1* (x-axis) and *POU4F1* (y-axis) in each cell from cluster 14. Each dot is an individual cell. Each cell is colored by its time course sample. **l)** wnnUMAP highlighting cells expressing exclusively *MYOD1* (pink), exclusively *PRDM12* (orange), or both (yellow). **m)** Scatter plot showing expression of *MYOD1* (x-axis) and *PRDM12* (y-axis) in each cell from cluster 6. Each dot is an individual cell. Each cell is colored by its time course sample. **n)** Scatter plot showing expression of *MYOD1* (x-axis) and *PRDM12* (y-axis) in each cell from cluster 14. Each dot is an individual cell. Each cell is colored by its time course sample. **o)** Quantification of iARMS cells that are HA (P3F)-expressing (gray), MYOD1-expressing (magenta), POU4F1-expressing (yellow), MYOD1-plus POU4F1-expressing (purple), and POU4F1-plus MYOD1-expressing (green). Data represented as mean +/- SEM. N = 5 images.

**Supplemental Figure 5.**
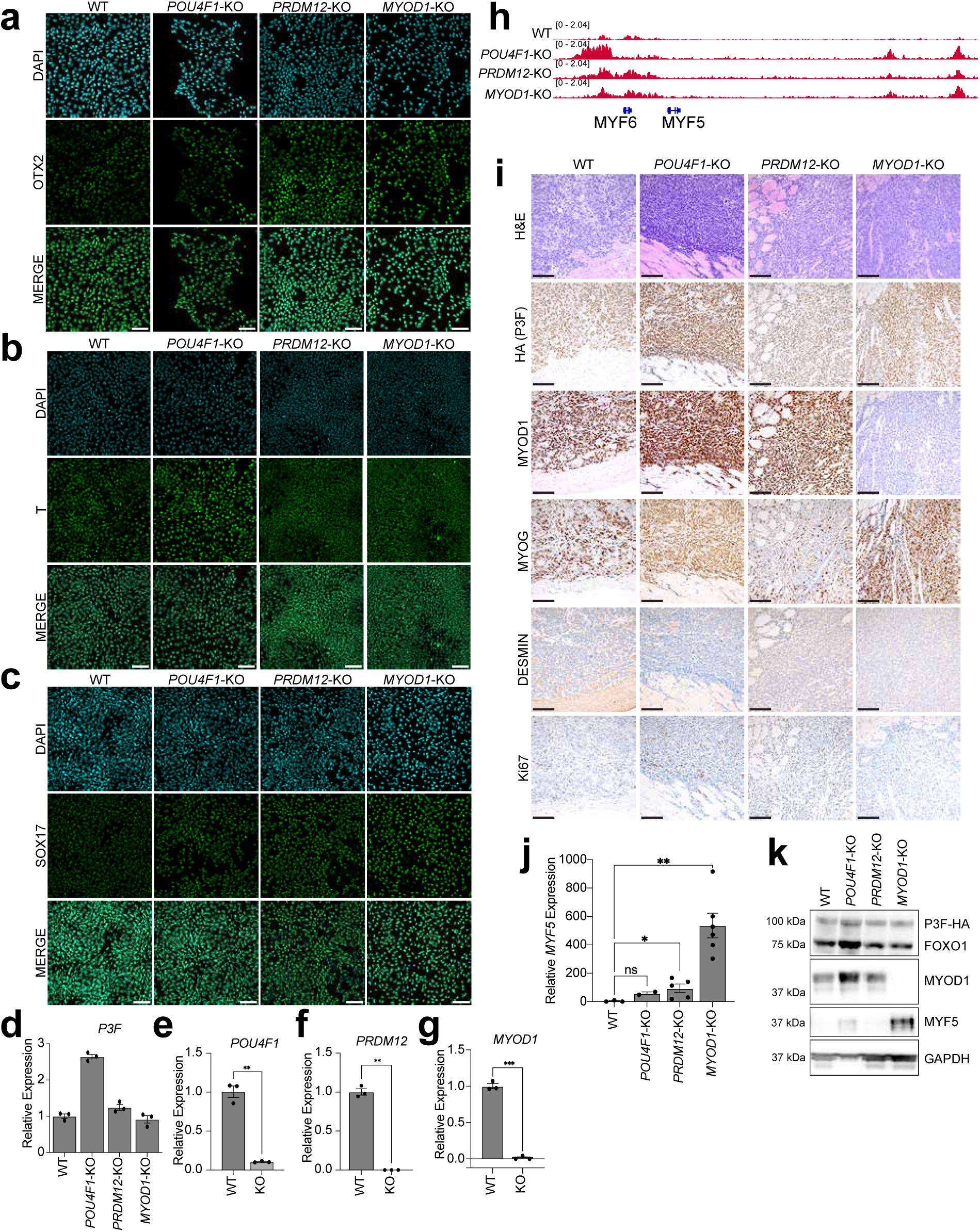
TF-KO iARMS cells and xenografts have altered myogenic gene expression. **a)** Representative immunofluorescence of *MYOD1*-KO, *PRDM12*-KO, *POU4F1*-KO, and WT BJFF.6 iPSCs for DAPI (cyan) and ectoderm marker OTX2 (green). Scale bar = 100 µm. **b)** Representative immunofluorescence of *MYOD1*-KO, *PRDM12*-KO, *POU4F1*-KO, and WT BJFF.6 iPSCs for DAPI (cyan) and mesoderm marker T (green). Scale bar = 100 µm. **c)** Representative immunofluorescence of *MYOD1*-KO, *PRDM12*-KO, *POU4F1*-KO, and WT BJFF.6 iPSCs for DAPI (cyan) and endoderm marker SOX17 (green). Scale bar = 100 µm. **d)** *PAX3::FOXO1* expression by real time PCR for WT, *POU4F1*-KO, *PRDM12*-KO, and *MYOD1*-KO iARMS. n = 3 technical replicates. **e)** *POU4F1* expression by real time PCR for WT and *POU4F1*-KO iARMS cells. n = 3 technical replicates. **f)** *PRDM12* expression by real time PCR for WT and *PRDM12*-KO iARMS cells. n = 3 technical replicates. **g)** *MYOD1* expression by real time PCR for WT and *MYOD1*-KO iARMS cells. n = 3 technical replicates. **h)** CUT&RUN alignment tracks for H3K27ac in WT, *POU4F1*-KO, *PRDM12*-KO, and *MYOD1*-KO iARMS cells at the gene locus for *MYF5* and *MYF6*. **i)** Representative histology (H&E) and immunohistochemistry (HA (P3F), MYOD1, MYOG, DESMIN, and Ki67) of WT, *POU4F1*-KO, *PRDM12*-KO, and *MYOD1*-KO iARMS tumors. Scale bar = 100 µm. **j)** *MYF5* gene expression by real time PCR for WT (n = 3 tumors), *POU4F1*-KO (n = 2 tumors), *PRDM12*-KO (n = 5 tumors), and *MYOD1*-KO (n = 6 tumors) iARMS xenografts. **k)** Immunoblots of PAX3::FOXO1, MYOD1, MYF5, and corresponding GAPDH in representative WT, *POU4F1*-KO, *PRDM12*-KO, and *MYOD1*-KO iARMS xenografts. All p-values in pairwise comparisons were determined by Student’s t-test (unpaired, two-tailed); *p < 0.05, **p < 0.01, ***p < 0.001, ****p < 0.0001. Data represented as mean +/- SEM.

## Methods

### Animals

All mouse strains except *Sox8^flox^* have been previously reported. Strains include *Myf6-Cre*^53^ (NCI 01XBL), *Pax3^Pax3::Foxo1^* ^41^(NCI 01XBM), *Trp53^flox^* ^54^(The Jackson Laboratory #8462), *R26-tdTomato* ^55^(The Jackson Laboratory #7914), and *NSG* (The Jackson Laboratory #005557). All tumor mice were maintained in mixed genetic backgrounds, so littermate controls were used for comparisons. For Kaplan-Meier survival analysis, all mice were observed from birth and sacrificed at humane endpoints, such as signs of obvious tumor or other distress. The maximum tumor burden/size permitted by the SJCRH Institutional Animal Care and Use Committee (IACUC) is 2 cm^3^. This burden was not exceeded in this study. All mice had consistent access to food and water. All mice were housed at ambient temperature (20-25 °C) and humidity (40-60%) with 12-hour light/dark cycles. All animal experiments were reviewed and approved by the SJCRH IACUC.

### Generation of *Sox8-flox* genetically engineered mouse

The *Sox8-flox* mouse model was created using CRISPR technology and direct embryo injection in the Center for Advanced Genome Engineering (CAGE, SJCRH, RRID:SCR_027027) and the Genetically Engineered Mouse Models Shared Resource (SJCRH). Briefly, 3-4-week-old C57BL/6N female mice from Jackson Labs (The Jackson Laboratory #005304) were superovulated with 5 units of gonadotrophin from pregnant mare’s serum (ProSpec Protein Specialists) and 48 hours later, with 5 units of human chorionic gonadotrophin (MilliporeSigma). After overnight mating with C57BL/6J males, the females were euthanized, and oocytes were harvested from the ampullae. A mixture of the sgRNAs, Cas9, and ssODNs consisting of 60ng/ul of Cas9 protein (St. Jude Protein Production Core, SJCRH), 20ng/ul of each sgRNA (Synthego), and 10ng/ul of each ssODN (IDT) were injected into the pronucleus of oocytes. The injected oocytes were then returned to culture media (M16 from Millipore or A-KSOM from Millipore) and later the same day transferred to day 0.5 pseudopregnant fosters (7–10-week-old CD-1 females from Charles River Laboratories mated to vasectomized CD-1 males). Resulting pups were genotyped via targeted amplicon sequencing as previously described^56^. Animals positive for both LoxP sites were backcrossed to C57BL/6J mice, tested to confirm that the LoxP sites were in cis, and then bred to homozygosity.

### Xenografts

All xenografts were performed in NSG mice. For ddP3F xenograft studies, NSG mice were maintained on doxycycline water at a concentration of 25 mg/kg. Cells were detached from plates with Accutase (Stem Cell Technologies 07922), resuspended in appropriate growth media, and counted. Cells were then pelleted by centrifugation at 100 x g and resuspended in 50% Matrigel basement membrane matrix (Corning 354234) diluted with appropriate growth media at a concentration of 5 million cells per 100 µL. This cell solution was kept on ice until injection. NSG mice were injected with 100 µL of cell solution into the gastrocnemius muscle using an insulin syringe. Both male and female mice were used as xenograft recipients.

### Cell Lines

293T (female origin, SJCRH, Martine Roussel), Rh30 (male origin, ATCC CRL-2061), Rh41 (female origin, SJCRH, Gerard Grosveld), Rh4 (female origin, SJCRH, Gerard Grosveld), JR (female origin, SJCRH, Adam Durbin), and Rh3 (male origin, SJCRH, Gerard Grosveld) cells were maintained in DMEM (HyClone SH32043) supplemented with 10% fetal bovine serum (HyClone SH30910) and 1% antibiotic/antimycotic (Sigma A5955). LHCN-M2 (male origin, University of Texas Southwestern, Woodring Wright) cells were maintained in DMEM/Medium 199 (Gibco 11965-084 and Gibco 11150-059) supplemented with 15% fetal bovine serum (HyClone SH30910), 0.02M HEPES (Gibco 15630), 0.03 µg/mL zinc sulfate (Sigma Z4750), 1.4 µg/mL Vitamin B12 (Sigma V2876), 0.055 µg/mL Dexamethasone (Sigma D-2915), 2.5 ng/mL hepatocyte growth factor (Chemicon Int’l GF116), and 10 ng/mL basic fibroblast growth factor (PeproTech AF-100-18B) on 0.1% gelatin (Sigma G1890-500G) coated dishes^57^. TP53 WT and KO BJFF.6 iPSCs (male origin, SJCRH, Shondra Pruett-Miller) as well as AN1.1 iPSCs (female origin, SJCRH, Shondra Pruett-Miller) were maintained in mTeSR Plus (Stem Cell Technologies 100-0276) and grown on either growth-factor reduced Matrigel basement membrane matrix (Corning 354230) or hESC-qualified Matrigel basement membrane matrix (Corning 354277). HUVECs (origin sex pooled donors, LifeLine Cell Technologies, FC-0044) were maintained in VascuLife VEGF Endothelial Medium Complete Kit (LifeLine Cell Technologies, LL-0003). All iARMS lines were maintained in “complete EGM” comprised of EGM BulletKit (Lonza cc-3124) supplemented with 20 ng/mL VEGF (PeproTech 100-20), 20 ng/mL bFGF (PeproTech AF-100-18B), and 1 µM CHIR-99021 (Stem Cell Technologies 72054). TP53 KO ddP3F “P3F on” cells were maintained in complete EGM supplemented with 10 µg/mL doxycycline hyclate (ThermoFisher J60579.14). All iARMS and ddP3F cells were cultured on 0.1% gelatin (Sigma G1890-500G) coated plates. All cells were maintained in a humidified incubator at 37 °C and 5% CO_2_.

### Molecular cloning and viral transduction

The lentiviral vector SJL21i TRE-Ch-M-Pur3G, was engineered to contain two separate expression cassettes in reverse orientation to the vector genome. The first cassette encodes the mCherry fluorescent reporter protein and is expressed by a third-generation tetracycline responsive promoter (TREG3G, Takara Bio, Mountain View, CA). The second cassette, controlled by a retroviral MND promoter, encodes a bicistronic open reading frame expressing puromycin N-acetyltransferase and the Tet-On 3G transactivator protein (Takara Bio) separated by a porcine teschovirus-1 2A self-cleaving peptide (P2A). This vector plasmid was constructed using a two-fragment ligation strategy into the lentiviral vector plasmid SJL12-MND-GFP digested with KpnI and MluI. The first fragment, obtained from SJL224v2 after digestion with KpnI and AgeI, contained the TREG3G promoter and mCherry sequences. The second fragment, derived from GFP-Dox-MND-puro using AgeI and MluI, included the puromycin N-acetyltransferase, P2A peptide, and the Tet-On 3G transactivator protein open reading frames. To make SJL21i TRE-Ch-E-Pur3G, the MND promoter was excised with EcoRI and AgeI, and a PCR amplified EF1a short promoter fragment engineered with EcoRI and BsmBI restriction sites was cloned into the plasmid to make SJL21i TRE-Ch-E-Pur3G. The EF1a promoter was amplified from SJL12-EF1a-GFP using the following primers: EF1 FW 5’-AGCTTGAATTCTGGCTCCGGTGCCCG and EF1 REV 5’-CTAGCTCCGTCTCACCGGTTCCTCACGACACCTGTGTTCTGG.

### Generation of TF KO BJFF.6 iPSCs

Knockout BJFF.6 iPSCs were generated using CRISPR technology at the Center for Advanced Genome Engineering (CAGE, SJCRH, RRID:SCR_027027). Briefly, 1 × 10 BJFF.6 cells (Cellosaurus, CVCL-VU02) were nucleofected using the Lonza 4D-Nucleofector™ X Unit with prel⍰assembled ribonucleoprotein complexes containing 100 pmol of each chemically modified sgRNA (IDT), 50 pmol of 3×NLS *Sp*Cas9 protein (St. Jude Protein Production Core), and, when applicable, 100 pmol of ssODN donor (IDT, Altl⍰R modifications). Nucleofection was performed in 20l⍰µL cuvettes using Solution P3 and program CAl⍰137 following the manufacturer’s instructions. Cells were single cell sorted five days post nucleofection on viability at the Flow Cytometry and Cell Sorting Shared Resource (St. Jude) and plated in prewarmed (37 °C) StemFlex (Thermo Fisher Scientific) supplemented with 5% CloneR2 (Stem Cell Technologies) onto Vitronectin XF (Stem Cell Technologies) coated 96-well plates. Clones were screened for the desired modification via PCR assays and targeted deep sequencing on a Miseq Illumina sequencer as previously described^56^. NGS analysis of clones was performed using CRIS.py^58^. Correctly modified clones were expanded and sequence confirmed. Cell line identity was authenticated using the PowerPlex® Fusion System (Promega) performed at the Hartwell Center (SJCRH) and tested negative for mycoplasma by the MycoAlert^TM^Plus Mycoplasma Detection Kit (Lonza). Editing construct sequences and relevant primers are listed in Supplemental Table 1.

### Histology and immunohistochemistry

For histologic analysis and immunohistochemistry, formalin-fixed paraffin-embedded (FFPE) sections were prepared by first fixing tissues in 10% neutral-buffered formaldehyde (NBF, Scigen 1015) overnight and then paraffin embedding. Hematoxylin and eosin (H&E) and immunohistochemistry staining were performed using standard procedures. Antibodies and autostainer conditions are listed in Supplemental Table 2. Images were acquired using an Axio Scan.Z1 microscope (Zeiss).

### Immunofluorescence

For immunofluorescence and imaging, cells were rinsed twice with ice-cold PBS and then fixed with 4% paraformaldehyde diluted in PBS for 20 minutes at room temperature. Fixed cells were washed 3 times for 5 minutes each with 1% BSA in PBS (1% BSA, Millipore 12659). Fixed cells were then incubated in “perm-block” (1% BSA, 10% NDS (Abcam ab7475), 0.3% TritonX-100 (BioRad 161-0407)) for 45 minutes at room temperature. Then, cells were incubated in appropriate primary antibody cocktail diluted with perm-block overnight at 4C. Following overnight incubation, the cells were washed 3 times for 5 minutes each with 1% BSA. The cells were then incubated in appropriate fluorescent tagged secondary antibody cocktail diluted in 1% BSA for 1 hour at room temperature in the dark. Following this incubation, the cells were washed 3 times for 5 minutes each with 1% BSA. Finally, the cells were covered with PBS and imaged using a Leica DMi8 Thunder Imager inverted fluorescent microscope.

### Myogenic differentiation assay

For myogenic differentiation, cells were plated and allowed to reach 80-90% confluency. Then, cells were changed to DMEM (HyClone SH32043) supplemented with 2% horse serum and 1% antibiotic/antimycotic (Sigma A5955) and cultured for 5-7 days. Following 5-7 days of differentiation, the cells were then subjected to immunofluorescence as described above.

### Endothelial directed differentiation and iARMS generation

Endothelial directed differentiation was performed as done previously^17, 31, 59^. Briefly, iPSCs were directed to differentiate into hemogenic mesoderm by first culturing in mTeSR Plus (Stem Cell Technologies 100-0276) supplement with 1 µM CHIR-99021 (Stem Cell Technologies 72054) and 10 µM ROCKi (Y-27632, Stem Cell Technologies 72302). The following day, the differentiating cells were changed to RPMI 1640 (ThermoFisher 11875093) supplemented with 50 ng/mL activin A (R&D Systems 338-AC) and 1X B-27 without insulin (Life Technologies A18956-01). The next day, the differentiating cells were changed to RPMI 1640 (ThermoFisher 11875093) supplemented with 40 ng/mL BMP4 (R&D Systems 314-BP), 1 µM CHIR-99021 (Stem Cell Technologies 72054), and 1X B-27 without insulin (Life Technologies A18956-01). Seventeen hours later, the cells were changed to StemPro-34 (Life Technologies 10639-011) supplemented with 4 mM 1-Thioglycerol (Sigma M6145), 2mM L-glutamine (Life Technologies 25030-081), 50ug/mL L-Ascorbic acid 2-phosphate sesquimagnesium salt hydrate (Sigma A8960-5G), 10 ng/mL BMP4 (R&D Systems 314-BP), 5 ng/mL bFGF (PeproTech AF-100-18B), and 300 ng/mL VEGF (PeproTech 100-20) and left for 72 hours. The end of this 72-hour incubation is designated as Day 5 in the iARMS generation time course. Untransduced endothelial cells were generated by culturing Day 5 cells in complete EGM (see “Cell Lines” for recipe) until Day 15, changing media every other day. iARMS cells were generated by transducing Day 5 cells with a lentiviral vector for P3F in complete EGM supplemented with 8ug/mL polybrene (Sigma TR-1003) and 10 µM ROCKi (Y-27632, Stem Cell Technologies 72302). Lentiviral media was changed on transduced cells 18 hours later, and subsequent media changes were performed every other day thereafter. ddP3F “P3F on” cells were generated by culturing Day 5 cells in complete EGM supplemented with 10 µg/mL doxycycline hyclate (ThermoFisher J60579.14) until Day 15 with media changes occurring every other day during the time course.

### iPSC germ layer differentiation

The Human Pluripotent Stem Cell Functional Identification Kit was used to validate pluripotency in iPSC models (R&D Systems SC027B). The kit was followed according to manufacturer’s instructions.

### Immunoblotting

SDS-PAGE and immunoblotting were performed using standard techniques as previously described. Briefly, protein lysates were prepared using RIPA lysis buffer supplemented with 1X PPIC (Roche 11697498001 and Roche 4906845001). Protein concentration was determined using BCA quantification assay (Pierce 23225) and equally loaded lysates were separated using SDS-PAGE. Transferred blots were incubated in primary antibody overnight at 4C. Blots were then washed and incubated with species specific secondary antibodies that were conjugated to horseradish peroxidase. Protein bands were visualized using either luminol reagent (Santa Cruz sc-2048) or Clarity Max Western ECL Substrate (BioRad 1705062) on an Azure 600 imager (Azure Biosystems).

### RNA isolation and gene expression analysis

Total RNA was isolated from samples using the miRNeasy micro kit (Qiagen 217084) according to the kit instructions. cDNA was synthesized from extracted RNA using first stand synthesis by Superscript III First Strand Synthesis using random hexamer primers (ThermoFisher 18080051). Quantitative real-time PCR was performed utilizing SYBR primers or TaqMan probes and normalized to β-actin RNA. Relative expression of genes was determined using ΔΔCT compared to β-actin. RNA sequencing was performed on purified RNA using the Illumina NovaSeq6000 genome sequencing system. RNA sequencing libraries were prepared using Illumina total stranded kit following manufacturer’s instructions (Illumina 20040525). Paired end sequencing was used for 100 base pair reads.

### Bulk RNA-sequencing analysis

Raw FASTQ sequences were aligned using HISAT2 (version 2.1.0) in paired-end mode using default parameters to version hg38 of the human genome to which the PAX3::FOXO1-HA sequence and the sequences of the External RNA Controls Consortium (ERCC) synthetic spike-in RNAs had been added as extra pseudochromosomes (Supplemental Table 3, http://tools.invitrogen.com/downloads/ERCC92.fa)^60^. Expression per-gene was quantified using htseq-count with parameters “htseq-count -i gene_id --stranded=reverse -f bam -m intersection-strict,” and gene annotation for GRCh38.p13 (GCF_000001405.39; Annotation Release 109) from RefSeq to which ERCC coordinates and PAX3::FOXO1-HA pseudo-coordinates were added^61^. We used per-gene read counts as input for differential expression analysis by DESeq2^62^. For differential expression analysis, raw count data were used as input to DESeq2^62^. Size factors were estimated using ERCC spike-in transcripts, followed by standard dispersion estimation and negative binomial modeling from DESeq2. Significantly differentially expressed genes were defined as those with |log_2_ fold change| ≥ 1 and adjusted p-value ≤ 0.05.

For certain displays, TPM (Transcripts Per Million) normalized expression was computed for each gene and spike-in probe. The standard TPM-normalization strategy is: normterm = sum of (readcount * readlength / exonlength) across all genes. TPM = readcount * readlength / exonlength * 1e6 / normterm. Exon lengths per-gene were calculated by collapsing all exons from all transcripts with the same gene name into a single set using BEDtools (v2.25.0) merge^63^. We normalized per-gene TPM values using the identically calculated ERCC spike-in values and a previously published analysis strategy^64^. Briefly, for each gene for each sample, we set the minimum expression value to 0.01 and add a pseudocount of 0.1. Using the affy R package, we used normalize.loess with the ERCC spike-in values of the expression table as the fitting subset.

### Gene Set Enrichment Analysis (GSEA) for Bulk RNA-seq

We performed Gene Set Enrichment Analysis (GSEA) using a preranked approach for analyses not based on direct condition-to-condition comparisons for “P3F on”, “P3F off”, and “P3F off high pass” ddP3F cells^65^. Genes were ranked by expression (TPM) rather than differential expression. For conditions with biological replicates, TPM values were averaged prior to ranking. To reduce noise, only genes within the top 50% by mean expression (TPM) were retained. Enrichment analysis was conducted using the parameters and gene sets as described below.

We performed GSEA using a preranked approach based on differential expression analysis on time-course data. Genes were ranked by normalized Log_2_ fold changes (LFCs) derived from DESeq2. To reduce noise, only genes within the top 50% by mean normalized expression (baseMean) were retained. Enrichment analysis was conducted using the fgsea and clusterProfiler packages in R. Enrichment analysis was conducted using the same parameters and gene sets as described below.

For bulk RNA-seq TF-KO samples which are lacking biological replicates, ERCC spike-ins were used to estimate size factors for normalization of raw counts. LFCs were then derived from the ERCC-normalized counts and used to rank genes for preranked GSEA. To reduce noise, only genes within the top 50% by mean normalized expression counts were retained. Enrichment analysis was conducted using the fgsea and clusterProfiler packages in R, using the same parameters and gene sets as described below.

GSEA analysis utilized the following specific parameters:

TERM2GENE: Our input gene set list combined the gene sets from “h.all.v2023.1.Hs.symbols.gmt” gene sets, as well as the ZHANG_DIRECT_P3F_TARGETS, REN_ALVEOLAR_RHABDOMYOSARCOMA_UP, REN_ALVEOLAR_RHABDOMYOSARCOMA_DN, DANIELLI_PROLIFERATIVE, DANIELLI_NEURONAL, DANIELLI_PROGENITOR, DANIELLI_DIFFERENTIATED, DAVICIONI_MOLECULAR_ARMS_VS_ERMS_UP, DAVICIONI_MOLECULAR_ARMS_VS_ERMS_DN, DAVICIONI_TARGETS_OF_PAX_FOXO1_FUSIONS_DN, EBAUER_TARGETS_OF_PAX3_FOXO1_FUSION_DN, GRYDER_PAX3FOXO1_TOP_ENHANCERS, DAVICIONI_RHABDOMYOSARCOMA_PAX_FOXO1_FUSION_UP, DAVICIONI_TARGETS_OF_PAX_FOXO1_FUSIONS_UP, TARGETS_OF_P3F_MELTZER_CHIPSEQ, DESCARTES_MAIN_FETAL_VASCULAR_ENDOTHELIAL_CELLS, DESCARTES_FETAL_MUSCLE_VASCULAR_ENDOTHELIAL_CELLS, DESCARTES_MAIN_FETAL_LYMPHATIC_ENDOTHELIAL_CELLS, and ENDOTHELIAL_CELLS_PANGLAO gene sets to align with the study’s focus on particular cellular phenotypes^14, 15, 23, 35, 38, 66–69^.

eps: Set to zero to ensure computational stability and prevent any potential division by zero errors.

nPermSimple: 10,000 permutations

pvalueCutoff: reporting p value cutoff of 1.

The results were visualized using the “ggplot2” and “pheatmap” package in R.

### Nuclei isolation and cleavage under targets and release using nuclease (CUT&RUN)

Nuclei were isolated by incubating cells in Nuclear Extraction Buffer (NEB) consisting of 20□mM HEPES KOH pH7.9, 10 mM KCl, 0.1% Triton X-100, 20% Glycerol, EDTA-free Protease Inhibitor (Roche 11836170001) for 10 minutes on ice. Aliquots of 500,000 nuclei in 100 µL of NEB were slowly frozen at -80C using a Mr. Frosty freezing container (Sigma C1562).

CUT&RUN was performed using the CUTANA ChIC/CUT&RUN kit (EpiCypher 14-1048) following manufacturer instructions and using 500,000 nuclei and 1 µg of antibody per reaction. CUT&RUN sequencing libraries were made using the CUTANA CUT&RUN Library Prep Kit (Epicypher 14-1001 and 14-1002) following manufacturer instructions. CUT&RUN libraries were sequenced using the NovaSeq6000 platform (Illumina). Paired end sequencing was done for 100 base pair reads.

### CUT&RUN Analysis

For CUT&RUN data, the first 2 bp of each read were uniformly trimmed across all samples and batches due to consistent sequence quality issues, and the trimmed reads were initially aligned to the E. coli reference genome (Escherichia coli K-12 MG1655) using Bowtie v1.2.2 in paired-end mode with the parameters -k 1 --best --un to retain unmapped reads, and -l set to the read length^70^. The remaining non-E. coli reads were then mapped to the human reference genome (hg38) using Bowtie with the parameters -k 2 -m 2 --best, and -l set to the read length. Transformations of the mapped read files, including Samtools sort -n, BEDtools bamToBed - bedpe, a manual conversion from BEDPE to BED3, and BEDtools bedToBam, were employed to generate BAM files containing CUT&RUN fragments from each aligned read-pair^63, 71^. For visualization purposes, the reference genome was partitioned into 50 bp windows using BEDtools makewindows, and fragment coverage across these windows was calculated using BEDtools intersect -c. The coverage was normalized per million mapped fragments, then converted into bedGraph format using bedGraphToBigWig, further into wiggle format via bigWigToWig, and ultimately visualized in the IGV (version 2.16.2) browser for human samples aligned to hg38^72, 73^. Because E. coli spike-in was not uniformly added across samples, spike-in–based normalization was not performed.

To identify regions of significant enrichment in CUT&RUN reads, MACS1.4 was employed^74^. Prior to peak-calling, individual reads from each paired-end alignment were discarded if they overlapped with the ENCODE-defined Problematic Regions list for the GRCh38 genome were excluded using BEDTools intersect with the -v option^75^. The remaining reads were used for peak calling against each sample’s corresponding IgG control, which was processed identically. The parameters --keep-dup=auto and -p 1e-9 were specified. To assess differences in coverage at a uniform set of regions of interest, peaks identified across samples were collapsed by using BEDTools merge. To separate enhancers from promoter-proximal regions, promoter regions were defined as ±2 kb from transcription start sites (TSS) based on the canonical GRCh38 gene annotation, and merged peaks overlapping these promoter regions were excluded using BEDTools intersect with the -v option. Remaining peaks were considered enhancers and were annotated to the gene with the nearest transcription start site using BEDTools closest with the -t first option and a BED file with the TSS of each gene. Read coverage for each sample within these enhancer regions was quantified using BEDTools intersect -c, and counts were normalized to reads per million mapped fragments (RPM) for comparative analysis. Log_2_ fold-changes in coverage were calculated after adding a pseudocount of 1.

### Single cell multiome (GEX + ATAC)

To perform single cell multiome, firstly, nuclei were isolated following 10X nuclei isolation guidelines (10X Genomics). 16,000 nuclei were used per lane of on the 10X Chromium iX platform. Samples were processed and sequencing libraries were prepared following manufacturer protocol (10X Genomics). Libraries were sequenced with the NovaSeq6000 (Illumina) platform following 10X sequencing guidelines.

### Preparation of Customized Reference Genome for Cell Ranger ARC Analysis

To enable detection of reads originating from the PAX3-FOXO1-HA lentiviral vector, a custom reference genome was built based on the GRCh38 (2020-A) assembly. Starting from the standard 10x Genomics GRCh38-2020-A reference package, we manually added a synthetic chromosome (chr24) containing the full mCherry-LTR cassette sequence (Supplemental Table 3). The corresponding gene model was appended to the GTF file by defining a new transcript (mcherryLTR_000001, Supplemental Table 3) with one exon spanning the full inserted sequence. This modified FASTA and GTF were used to rebuild the 10x Genomics reference genome using **Cell Ranger ARC v2.0.2** (https://www.10xgenomics.com/support/software/cell-ranger-arc/latest) with the command: cellranger-arc mkref --config=“GRCh38-2020-A-build/P3F_hg38.config”. The resulting custom reference, named GRCh38_mcherryLTR, includes all standard GRCh38 chromosomes plus chr24 for the reporter cassette.

### Cell Ranger ARC Data Processing

Single-cell multiome (ATAC-seq + gene expression) libraries were generated at multiple experimental timepoints: Day 0 (iPSC) and Day 5 (immediately prior to P3F transduction), and Days 7, 9, 11, and 15 under both P3F-positive and UT conditions.

Raw sequencing data were processed using **Cell Ranger ARC v2.0.2** software with parameters --localcores=16, --localmem=64 (https://www.10xgenomics.com/support/software/cell-ranger-arc). Each dataset was analyzed independently using the custom reference described above. Each run produced paired gene-expression and chromatin-accessibility count matrices aligned to the extended GRCh38_mcherryLTR genome.

### Multiome Data Integration and Merged Seurat Object Construction

Filtered multiome (ATAC-seq and gene expression) outputs generated by Cell Ranger ARC (v2.0.2, 10x Genomics) were used as input, including filtered feature-barcode matrices (filtered_feature_bc_matrix.h5) and ATAC-seq fragment files (atac_fragments.tsv.gz), which retain barcodes identified as cell-associated by the pipeline. Data were imported into R (v4.4.1) and processed using Seurat (v4.0.3) and Signac (v1.12.0)^76, 77^. For each time point (Day 0–15; P3F transduced and UT conditions), paired RNA and ATAC assays were constructed. Gene annotations were obtained from the custom GRCh38_mCherryLTR reference generated during Cell Ranger ARC processing. Sample-specific Seurat objects were generated using CreateSeuratObject() and CreateChromatinAssay() without additional filtering at this stage, and subsequently merged across samples using merge(), with unique cell barcodes retained for each sample. To define a unified chromatin accessibility feature space, peak regions from all samples were combined using GenomicRanges::reduce(), retaining peaks with widths between 20 bp and 10 kb^78^. A shared ATAC-seq count matrix was then recomputed using FeatureMatrix() across all cells.

Quality-control metrics were calculated on the merged object, including the number of detected RNA features (nFeature_RNA), percentage of mitochondrial transcripts (percent.mt), number of detected ATAC features (nFeature_ATAC), transcription start site enrichment (TSS.enrichment), and nucleosome signal. Based on inspection of metric distributions, nuclei were retained if they met the following criteria: 300 < nFeature_RNA < 10,000, percent.mt < 30%, 1,000 < nFeature_ATAC < 40,000, TSS.enrichment > 1, and nucleosome signal < 7. The final filtered Seurat object containing integrated RNA-seq and ATAC-seq assays was used for downstream normalization and dimensionality reduction analyses.

### Batch effect normalization and dimensionality reduction

Batch effects across samples were mitigated using Cluster Similarity Spectrum (CSS) normalization implemented in the simspec package (https://github.com/quadbiolab/simspec)^79^. Cells were grouped by sample identity (orig.ident), and CSS embeddings were computed separately for RNA-seq and ATAC-seq modalities at a clustering resolution of 0.6. Cells with incomplete CSS embeddings were excluded from downstream analysis. For bimodal integration, Seurat’s weighted nearest neighbor (WNN) framework was applied using the CSS-corrected RNA and ATAC reductions (css_rna and css_atac). Multi-modal neighbors were identified with FindMultiModalNeighbors() using all available dimensions from each modality, and modality weights were computed for RNA-seq and ATAC-seq assays. A two-dimensional wnnUMAP embedding was generated from the weighted nearest neighbor graph (weighted.nn), and cell clustering was performed on the weighted shared nearest-neighbor graph (wsnn) using the Louvain algorithm at a resolution of 0.3. Clustering resolution was selected by evaluating multiple candidate resolutions and assessing cluster stability, separation in wnnUMAP space, and concordance with known marker gene expression patterns; a resolution of 0.3 was chosen as it provided well-resolved, biologically interpretable clusters without over-fragmentation.

### Single-cell pathway activity analysis using ssGSEA

Pathway activity in single cells was quantified using the single-sample gene set enrichment analysis (ssGSEA) method described by Barbie et al. 2009 and implemented in the GSVA package^80, 81^. In this context, each individual cell was treated as an independent sample for enrichment scoring. Custom gene sets were imported from the same pathway collection used for bulk RNA-seq preranked GSEA analysis described above and converted into a list format. Normalized gene expression values from the RNA-seq assay of the Seurat object (derived from single-cell multiome data) were used as input. A SingleCellExperiment object was constructed from the expression matrix, and only genes overlapping between the expression matrix and gene sets were retained. Enrichment scores were computed for each gene set in each cell based on the relative ranking of genes within that cell. The resulting scores were added to the Seurat object as cell-level metadata and visualized on the precomputed wnnUMAP embedding.

### Co-expression analysis

Expression patterns of selected marker genes were visualized across cell clusters using the DotPlot function in Seurat (v4.0.3). A curated set of genes, including ADAMTS5, PAX5, EBF3, MYOD1, POU4F1, C1QTNF3, PPP1R14C, PAX3, PRDM12, PITX3, DMRTA2, and the PAX3::FOXO1 reporter (mCherryLTR), was selected based on relevance to lineage identity and transcriptional programs^76^. Only genes detected in the RNA assay were included in the analysis. Cell identities were defined based on clustering results and ordered numerically. DotPlot visualizations display the percentage of cells expressing each gene within a cluster (dot size) and the average expression level (color intensity). For visualization purposes, genes were ordered based on their average expression within a representative cluster to highlight cluster-enriched expression patterns. Plots were generated using Seurat and ggplot2, with customized color scaling and axis formatting to facilitate comparison across clusters. Co-expression was visualized on wnnUMAP embeddings using the FeaturePlot function in Seurat with blended feature display enabled (blend = TRUE). Normalized RNA assay expression values were overlaid per cell, with color indicating relative single-gene expression and co-expression in the blended panel. Plots were further customized using ggplot2 and cowplot.

### Single-cell motif activity analysis

Motif activity was quantified from the ATAC-seq modality using chromVAR within the Signac framework. PFMs from the JASPAR2022 vertebrate CORE collection were retrieved, and a custom PAX3::FOXO1 fusion motif was manually defined, converted to a PFMatrix object using TFBSTools, and appended to the motif collection^82–84^. Motif annotations were added to the ATAC-seq assay using AddMotifs() with the reference genome sequence, and bias-corrected per-cell motif deviation scores were computed using RunChromVAR(). Motif activity for selected transcription factors was visualized on the wnnUMAP embedding, and sequence logos were generated using seqLogo (Bembom O, Ivanek R (2024). seqLogo: Sequence logos for DNA sequence alignments. R package version 1.70.0.)^85^.

### Dependency Analysis

Gene dependency scores were obtained from the DepMap resource (DepMap Public 25Q2 release). CRISPR-based gene dependency scores (Chronos) and RNAi-based dependency scores (DEMETER2) were used to assess dependency on PAX3 and FOXO1 across cancer cell lines. For each gene, cell lines were ranked based on their dependency scores, with lower scores indicating stronger dependency. Cell lines of interest were highlighted and annotated within the ranked distributions. Dependency score distributions were visualized by plotting ranked cell lines against their corresponding Chronos or DEMETER2 scores, with vertical reference lines indicating the score = 0. Comparisons between CRISPR-based (Chronos) and RNAi-based (DEMETER2) dependency estimates were performed to assess consistency of gene dependency patterns across platforms.

### Endothelial Tube formation assay

Matrigel (#354234, Corning) was defrosted overnight at 4C and 250 μL added to each well of a 12-well plate. After solidifying for 30 mins at 37C, 1 x 10^5^ HUVEC, ARMS, ddP3F “P3F on”, ddP3F “P3F off”, or LHCN cells were plated onto Matrigel in appropriate media and imaged every hour for 8 hours in an Incucyte S3 (Sartorius) or an Incucyte SX5 (Sartorius).

### Endothelial Tube formation analysis

Angiogenesis Analyzer for ImageJ was used to quantify the number and size of endothelial tubes. Incucyte images from the 8-hour timepoints were saved as .TIFF and batch uploaded into the Angiogenesis Analyzer tool and analyzed following standard protocol^86^. The number of meshes and the total mesh areas from each field of view were quantified in GraphPad Prism.

### Viability assay

Viability of cells was measured using the CellTiterGlo (Promega G7570) assay following manufacturer instructions. Luminescence readings were taken with a BioTek Synergy H1 plate reader (Agilent). Quadruplicate measurements were taken per condition.

### Proliferation assay

Proliferation measurements were collected using either an IncucyteS3 (Sartorius) or IncucyteSX5 (Sartorius). Briefly, cells were lifted and plated at a concentration of 10,000 cells per well of a 96-well plate in 200 µL of appropriate growth media. The cells were then placed in the Incucyte and tracked for 5-7 days with scans taking place every 4 hours and taking 4 images per well. Incucyte analysis software (Sartorius) was used to calculate confluency of each image and data was plotted as an average of 16 images per condition.

### Focus formation

For each focus formation assay, cells were lifted and plated at a concentration of 1,000 cells per well of a 6-well dish and cultured for 10 days. Cells were cultured in appropriate growth media and additional media was added every 3 days to replenish media lost to evaporation. After incubation for 10 days, the cells were rinsed with PBS and fixed in their respective wells with ice-cold methanol for 10 minutes. Foci were then stained with 0.02% crystal violet staining solution (0.2 g of crystal violet dissolved in 100 mL of a 2% ethanol solution) for 10 minutes at room temperature (Fisher Scientific C581-25). After staining, wells were rinsed twice with deionized water and allowed to air dry overnight before imaging. Plates were imaged using Epson scanner and images were prepared using ImageJ.

### Pluripotency assay

To test for pluripotency, iPSCs were subjected to the Human Pluripotent Stem Cell Functional Identification Kit (R&D Systems SC027B) following manufacturer instructions. Cells were fixed and immunofluorescence was performed as described above.

### Statistical analyses

Data analyses were performed using Microsoft Excel, GraphPad Prism (version 11.0.0), and the R software packages. Sample sizes and replicates are listed in relevant text and figure legends. All error bars are reported as mean ± SEM unless otherwise mentioned. Statistical significance was determined using Students t-test for pairwise comparisons unless otherwise mentioned (*P<0.05, **P < 0.01, ***P < 0.001, ****P < 0.0001).

## Data Availability of SJCRH Patient tumor RNA expression

SJCRH patient tumor RNA expression at diagnosis was obtained from St. Jude Cloud (https://www.stjude.cloud)^43^.

